# Concerted modulation of spontaneous behavior and time-integrated whole-brain neuronal activity by serotonin receptors

**DOI:** 10.1101/2024.08.02.606282

**Authors:** Drew Friedmann, Xavier Gonzalez, Ashley Moses, Tanner Watts, Anthony Degleris, Nicole Ticea, Jun H. Song, Sandeep Robert Datta, Scott W. Linderman, Liqun Luo

**Affiliations:** Department of Biology, Howard Hughes Medical Institute, Stanford University; Cajal Neuroscience Inc; Department of Statistics, Wu Tsai Neurosciences Institute, Stanford University; Department of Computer Science, University of Utah; Department of Electrical Engineering, Stanford University; Department of Physics, Stanford University; Department of Neurobiology, Harvard Medical School

**Author notes:** Corresponding authors. (SWL), (LL). Equal contribution (Drew Friedmann, Scott W. Linderman).

## Abstract

Serotonin neurons from the raphe nuclei project across the entire brain and modulate diverse physiology and behavior by acting on over a dozen receptors. Here, we took a step towards dissecting this complex process by examining the effects of agonists and antagonists of four widely expressed serotonin receptors (2A, 2C, 1A, and 1B) on spontaneous mouse behavior, which we related to time-integrated whole-brain neuronal activity as assessed by the expression of Fos, a canonical immediate-early gene product. Low-dimensional representations of behavioral and Fos map data revealed the dominant factors of variation in each domain, captured predictable differences across drug groups, and enabled predictions of behavioral changes following perturbations in Fos maps and vice versa. Our study provides a rich resource describing the effects of manipulating serotonin receptors on animal behavior and whole-brain integrated neuronal activity. It also establishes an experimental and analysis paradigm for interrogating the relationship between behavior and neuronal activity across different time scales.

## INTRODUCTION

A central goal of neuroscience is to link animal behavior with neuronal activity in the brain. This has traditionally been approached by focusing on a particular behavior and neuronal activity in a specific brain region. Much progress has been made from such focused studies, which have revealed principles underlying sensory perception, decision making, spatial navigation, and motor planning and action^1–4^. With recent technical advances, it is now possible to record or optically image activity of thousands of neurons simultaneously in behaving animals^5,6^. Substantial progress has also been made in methods that record and quantify the spontaneous behavior of animals without human supervision^7–10^. These advances create an opportunity for investigating the relationship between large-scale neuronal activity and natural behavior in the same animal.

Indeed, in simpler nervous systems with optical transparency, such as *C. elegans* and zebrafish larvae, researchers can perform optical imaging of most neurons in behaving animals^11–14^. However, measuring whole-brain activity with single-neuron resolution in a behaving mammal is not yet possible and will unlikely be in the foreseeable future. Large-scale optical imaging can only access neurons close to the surface of the skull, and high-density electrophysiological probes can only access but a tiny fraction of the many millions of neurons in a mammalian brain. An alternative to real-time neuronal activity is to measure time-integrated activity via immediate early gene (IEG) expression, which is rapidly and transiently induced by neuronal activity^15^. IEGs have been used to discover neuronal populations related to many specific behaviors and to manipulate neuronal populations that were previously activated by specific stimuli or behavioral tasks^16–19^. However, it is unclear the extent to which natural behavior relates to time-integrated activity in the whole brain.

Serotonin is a powerful neuromodulator of physiology and behavior in health and disease, and is therefore a natural candidate as an intervention to interrogate whole brain activity and behavior. In the mammalian brain, serotonin neurons reside in the raphe nuclei in the brainstem but project their axons across the entire brain^20,21^. Diverse functions have been ascribed to neuromodulation by serotonin, from sensory gating and locomotor activity to learning, social interactions, and regulation of diverse brain states such as anxiety and mood^22,23^. The diverse effects of serotonin on target neurons are mediated by 14 distinct serotonin receptors in mice that are expressed in specific brain regions and cell types^24–28^. Except for serotonin receptor 3, which is an ionotropic receptor, all serotonin receptors are metabotropic receptors that are coupled to specific G proteins, causing either depolarization (e.g., serotonin receptor 2 subfamily, coupled to G_q_/G_11_) or hyperpolarization (e.g., serotonin receptor 1 subfamily, coupled to G_i_/G_o_) of target neurons^25^. Because of medical relevance (e.g., in treating neuropsychiatric disorders), agonists and antagonists that preferentially target individual serotonin receptors have been extensively characterized^25,26^. Furthermore, investigating the effects of specific serotonin receptors on neural activity and animal behavior can also provide a powerful means to dissect the diverse roles played by the serotonin neuromodulatory system.

Here we used a panel of well-characterized serotonin receptor agonists and antagonists to investigate their effects on spontaneous behavior and whole-brain integrated activity of the same mice via the expression levels of Fos, an IEG product. We developed statistical models to probe the relationship between behavioral ‘syllables’ and Fos maps and established a platform of predicting changes in behavior following perturbations in Fos maps and vice versa. Our study provides a rich resource describing how spontaneous behavior and integrated whole-brain activity are modulated by activation and inactivation of specific serotonin receptors. It also establishes a paradigm with which to investigate the relationship between behavior and brain activity under specific conditions in an unbiased manner.

## RESULTS

We administered serotonin agonists/antagonists to 190 mice (9 drug groups and 1 saline control group, 19 mice per group) 30 min prior to monitoring their spontaneous behavior in a circular open field for 60 min. During the period between 30 and 45 min, a collimated white LED light illuminating the center of the arena was turned on, intending to heighten the anxiety of the mice. 45 min after behavior recording (to allow Fos expression), mice were sacrificed for whole-mount Fos immunostaining (**Fig. 1a**). We first present data and analyses on the effect of drugs on behavior (**Fig. 1** and **Fig. 2**). We then present data and analyses on the effect of drugs on whole-brain Fos levels (**Fig. 3–5**). We finally investigate the relationship between behavior and Fos maps (**Fig. 6**). The data described below came from the 188 mice that passed the quality control for behavioral data (**Fig. 1** only) and the 168 mice that passed the quality control of data for both behavior and Fos map (**Fig. 2–6**).

**Fig. 1.**
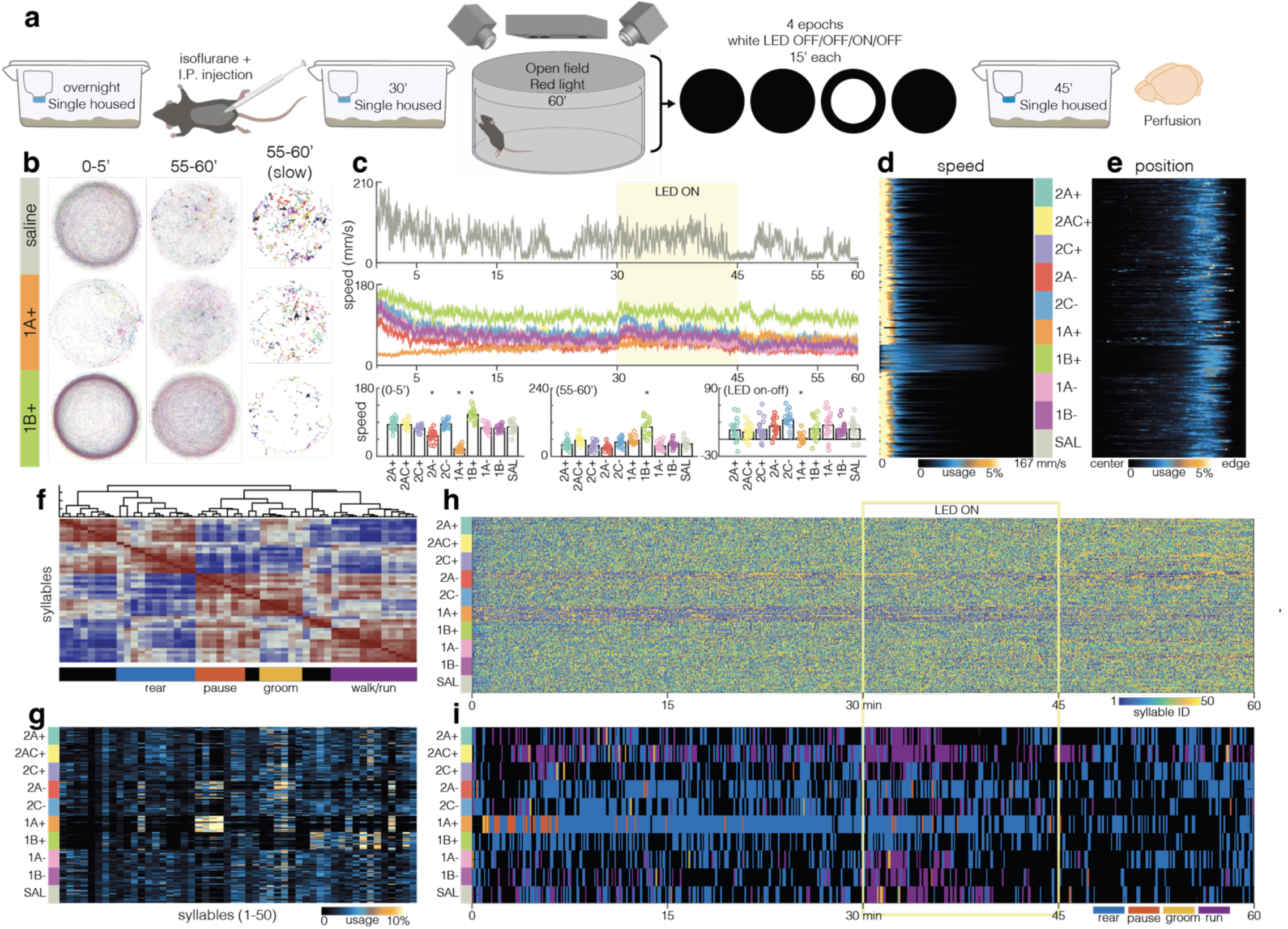
The effect of serotonin receptor agonists and antagonists on spontaneous behavior. **a**, Experimental design and timeline for drug dosing, open field behavioral recording (at 30 frames/sec), and histology for whole-brain Fos mapping. **b,** Overlaid trajectories of individual mice from 3 of the 10 drug conditions. All mice from these conditions (n = 19, 17, 19) are shown for the displayed time windows. The 3^rd^ column highlights trajectory positions with speed below 30 mm/s, highlighting that 1B+ reduces pauses. **c,** Top, smoothed 2D-speed recording for a single saline-treated mouse. The 15-min LED-on period is highlighted. Middle, average speed for each drug treatment group (color code as in **d**). The LED-on epoch is highlighted in yellow. Bottom (left and center), time binned average speed for each drug group during the first and last 5 min. Bottom right, the difference between the period 2 min after LED onset compared to 4 min prior. Group different from saline: * p < 0.05; ANOVA followed by Turkey post hoc test. **d,** Speed distribution for each mouse (rows), grouped by drug and measured as percentage of frame transitions representing each speed (mm/sec). **e,** Positional distribution for each mouse (rows), grouped by drug and measured as a percentage of frames spent at a given concentric position of the arena; 140 bins. **f,** Cladogram output of a spectral clustering of MoSeq syllables highlights groups of similar structure. Manual descriptions of blinded video from each syllable resulted in categorical descriptors for four major tree branches of the hierarchy. The remaining clusters grouped most closely with rearing and locomotion respectively but were less well described by users viewing video. **g,** Average usage (as a percentage of frames) of each syllable for all mice across drug groups aligned to the cladogram in **f**. **h,** Colormap of MoSeq syllables 1–50 shown for all mice (n = 188) and frames (n = 108000). Conditions 1A+ and 2A– stand out as horizontal bands, but all groups shift in syllable usage across time (blue to yellow). **i,** Differential usage of the syllables comprising four major categorical behaviors as described in **f** by drug group across the full recording session. Calculated as the modal category in rolling bins of 200 frames. The remaining clusters of syllables adjacent to rearing and locomotion are uncolored. LED response is seen as an increase in the walk/run syllables in the 2A+, 2AC+, 1A–, 1B–, and saline conditions.

**Fig. 2.**
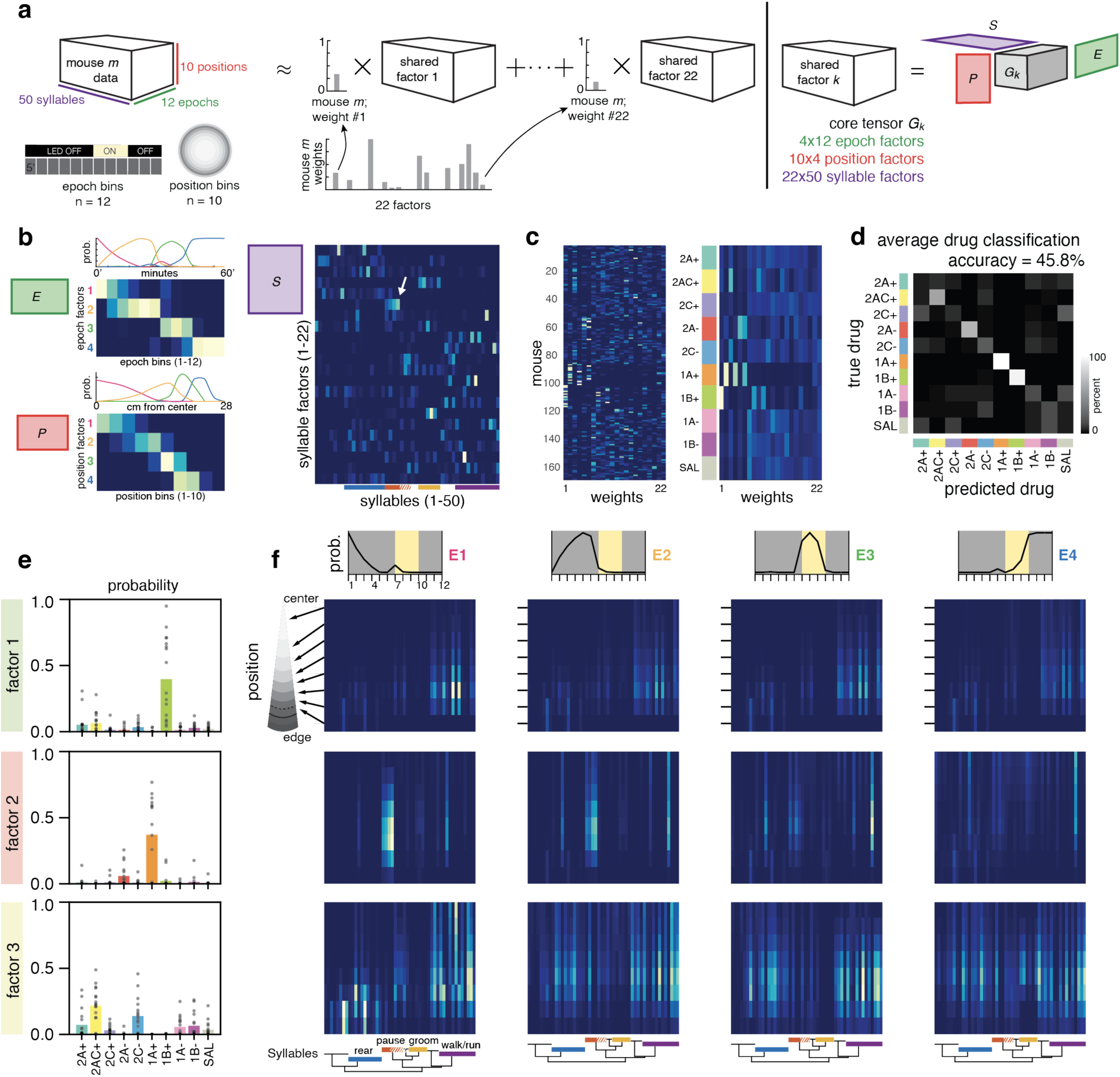
Decomposition of behavior. **a**, Graphical representation of the tensor-decomposition wherein each mouse’s behavioral recording is represented as a 3D tensor of frame counts for each combination of syllable (1–50), epoch (1–12; consecutive 5-min bins), and position (1–10; equal area rings, 1 being the center and 10 the periphery of the arena). The resultant factorization includes a shared core tensor *G*, one set of weights each for 4 epoch factors, 4 position factors, 22 syllable factors, and 22 mouse weights. **b,** Top left, the 4 epoch factors arrange into a cascade across the 12 five-min bins from start to end of the recording session. Bottom left, the 4 position factors arrange into a cascade across the 10 concentric position bins of equal area from the center to the edge/wall of the arena. Probability distributions (prob.) are shown above the heatmaps for epoch (*E*) and position (*P*). Right, the 22 syllable factors capture subsets of syllables with correlated usage. Subset of pausing syllables in factor 6 is indicated, white arrow. Color code along the lower axis represents rearing, pausing, grooming, and locomotion syllables as established in **Fig. 1**. **c,** The per-mouse weights for individual (left) and average for each drug group (right). **d,** Confusion matrix of drug classification based on tensor decomposition of behavior. The average accuracy is 45.8%, well above the baseline prediction is 10% given 10 groups. **e.** The weights on the first three factors differ substantially across drug groups. **f,** Factors contain information about distributions over what, when, and where syllables are used throughout the session. For example, factor 1 shows decreasing levels of locomotion, factor 2 shows elevated pausing, later transitioning to a specific form of locomotion, and factor 3 shows a transition from rearing and walking/running near the periphery to more diffuse behavior by the end of the session.

**Fig. 3.**
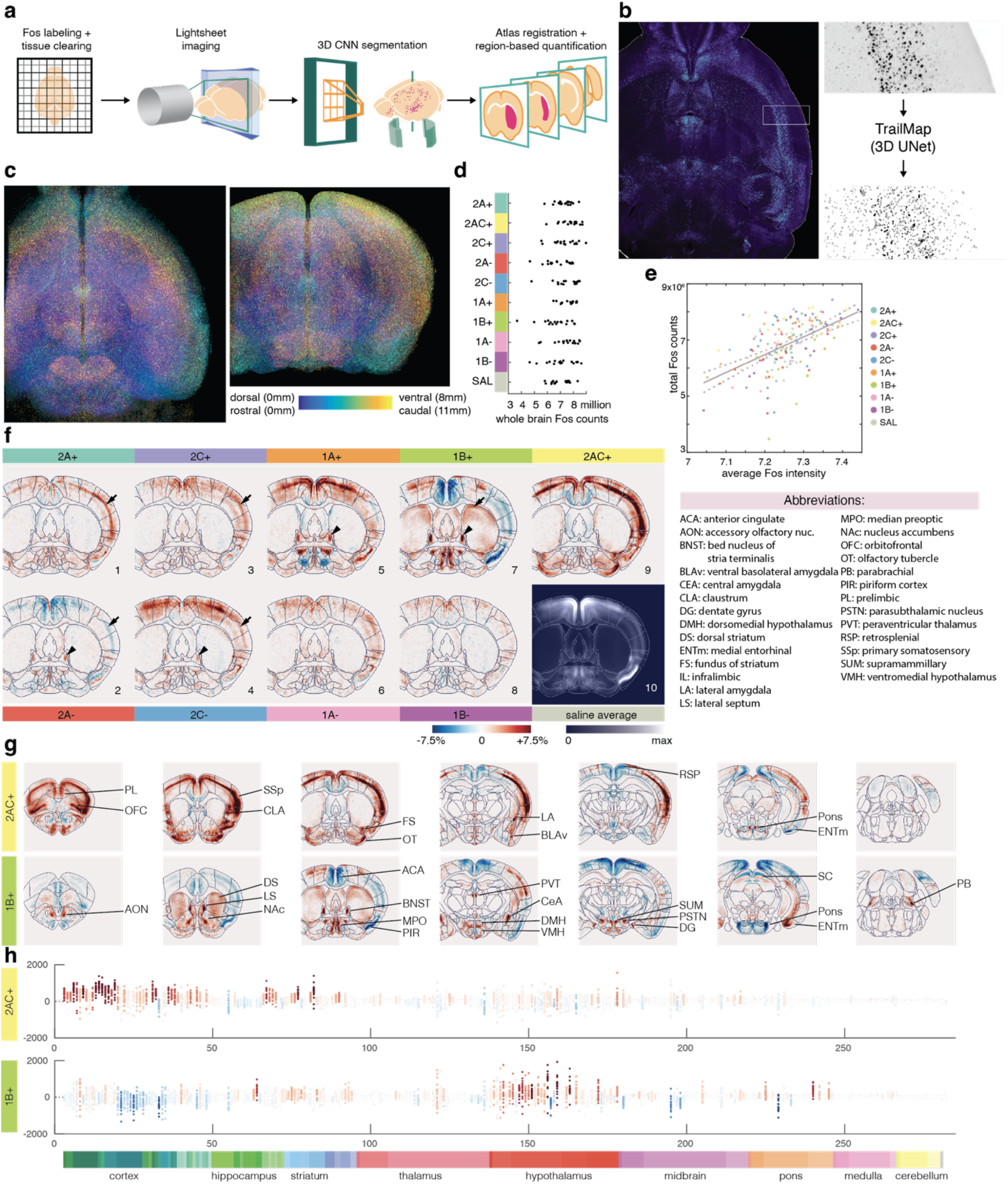
Overview of the effect of serotonin receptor agonists and antagonists on whole-brain Fos maps. **a**, Experimental design for whole-brain analysis pipeline of Fos image data. **b,** Left, horizontal view of a 500-µm Z-projection of raw Fos image data. Right, zoom-in of the boxed region from the left image (top), which is segmented by 3D-Unet based TrailMap. Centroids are established as local maxima detected from the probability maps and a 6-connected neighborhood of raw intensity values is averaged to be the intensity of each centroid. **c,** Horizontal (left) and coronal (right) depth-coded views of a single brain’s detected centroids demonstrate even coverage across the entire intact brain. **d,** Total numbers of counted nuclei per mouse and grouped by drug treatment. **e,** Scatter plot of average Fos intensity against the total Fos count (as in **d**) for all 168 mice, color coded by drug treatment. Linear fit shows significant correlation, R^2^ = 0.281, p = 1.4×10^-13^. **f,** A representative 100-µm coronal slice of saline-subtracted Fos intensity maps for each drug treatment group, averaged across individuals within treatment groups. **g,** Coronal slices (as in **f**) across anteroposterior axis for two example drug treatment groups. **h,** Average saline-subtracted Fos intensity for the two drug treatment groups in **g** for all nuclei in each of 282 Allen-brain regions at the bottom, color coded according to Allen Brain Atlas^45^. Each dot represents data from an individual mouse. Color represents the brain region’s average change from saline across all mice in the group (as in **f**).

Of the 14 serotonin receptors encoded by the mouse genome, we selected agonists and antagonists for their preferential activity on serotonin receptors 1A, 1B, 2A, and 2C (**Table 1**) because of their broad expression and functions in the brain^24–26^. For example, 1A receptors have been targeted in clinical trials for anxiety and depression^29^. 1B receptors are implicated in dopaminergic modulation, drug addiction, and aggression^30^. 2C receptors regulate food intake and body weight^31^. 2A agonists have seen a recent resurgence in both trials and research as they mediate the major effects of classical psychedelics^32^, which have shown promise in treatment of major depressive disorder. In the following text, we abbreviate the agonists as 1A+, 1B+, 2A+, and 2C+, and antagonists as 1A–, 1B–, 2A–, and 2B–, respectively. A 9^th^ drug, 2AC+, is a mixed agonist for both serotonin receptors 2A and 2C but has historically been a popular choice as a 2A agonist. Drug concentrations were chosen based on prior characterization of their pharmacological properties^33–40^.

**Table 1:** Serotonin receptor agonists and antagonists used in this study.

| Group<sup>1</sup> | Drug | Target<sup>2</sup> | Dose (mg/kg) |
| --- | --- | --- | --- |
| 1 | 25CN-NBOH | 2A+ | 3 |
| 2 | DOI | 2A/2C+ | 1 |
| 3 | PF-04479745 | 2C+ | 5 |
| 4 | MDL100907 | 2A– | 1 |
| 5 | SB-242084 | 2C– | 0.75 |
| 6 | NLX-101 | 1A+ | 0.75 |
| 7 | CP 94253 | 1B+ | 10 |
| 8 | (S)WAY-100135 | 1A– | 10 |
| 9 | SB-216641 | 1B– | 5 |
| 10 | saline | control | none |

<sup>1</sup> With the exception of Fig. 3f, we list drugs according to this sequence.
<sup>2</sup> Listed here are the serotonin receptors that have the highest affinity for a particular drug, even though most drugs have lower affinity binding to other receptors.

### Effects of drugs on spontaneous behavior

For control mice injected with saline, examination of their trajectories (**Fig. 1b**, **Extended Data** Fig. 1a) and speed (**Fig. 1c, d**) revealed that their highest locomotion activity occurred during the first 15 min (and particularly the first 5 min) in the circular arena, likely reflecting their exploration of this novel environment. Mice slowed down during the 2^nd^ 15-min period, increased locomotion in response to LED-on stimulation during the 3^rd^ 15-min period, and returned to a low-activity state in the final 15-min period. Most drug cohorts followed a similar trend. 1A+ treated mice (1A+ mice hereafter) were an exception. They exhibited reduced movement at the outset and did not significantly increase speed in response to LED-on (**Fig. 1c**, quantified in the bottom panels). By contrast, 1B+ mice exhibited increased locomotion throughout the observation period compared to control; they also showed an increase of locomotion in response to LED stimulation (**Fig. 1b–d**). Observation and quantitative analyses of spatial location revealed that all mice preferred to reside or move in the periphery, with 1B+ mice showing the strongest preference and 2C+ mice showing the largest shift to the periphery after LED onset (**Fig. 1b, e, Extended Data Fig. 1a, b**).

Unsupervised clustering of behavior using the motion sequencing (MoSeq) platform^7,10,41^ of all 188 mice during the 60-min period produced 50 “syllables,” which capture stereotyped movements lasting approximately 300 ms each (**Table S1, Extended Data** Fig. 2). Spectral clustering of the MoSeq syllables based on their kinematics produced hierarchical clusters that matched with independent human annotations of these syllables as “rear”, “pause”, “groom”, and “walk/run” (**Fig. 1f**). Comparing the frequency of syllable usage across different drug conditions revealed an increased usage of walk/run syllables and a decreased usage of pause syllables for 1B+ mice (**Fig. 1g**), consistent with increased locomotion speed from our scalar analysis above (**Fig. 1c**). By contrast, 1A+ mice exhibited a substantial increased usage of most pause syllables and decreased use usage of most walk/run syllables (**Fig. 1g**). Interestingly, 2A– mice exhibited similar syllable usage in pause and walk/run as 1A+ mice (**Fig. 1g**), suggesting shared behavioral effect of inhibiting 2A receptor and activating 1A receptor. As will be described later, 2A– and 1A+ also caused similar neuronal activation patterns in specific brain regions.

Drugs also affected syllable use over time. **Fig. 1h** presents usage distributions of all 50 syllables across the 60-min period for all mice. To simplify visualization of this complex dataset, we identified the most-frequently-used syllable cluster in rolling bins of 200 frames (∼6.7 seconds) across mice of each drug condition and displayed these across the temporal epochs (**Fig. 1i**). This analysis revealed that LED onset caused an increased use of run syllables in saline and most drug conditions. However, for mice treated with the 1A+ and 2A–, rearing appeared to be the dominant behavioral mode after LED onset (**Fig. 1i**).

### Decomposition of behavior

The trends in syllable usage, spatial occupancy, and running speed over time (**Fig. 1**) suggested that there are strong correlations in what, where, and when syllables are used by different drug groups. From a modeling perspective, such correlations would imply that the behavioral data is low dimensional. We developed novel methods to probe for low-dimensional, spatiotemporal patterns of syllable usage.

Typical MoSeq analyses consider egocentric pose and do not take absolute position or elapsed time into account^7,42^. To understand how syllable usage varied with position in the arena and the time elapsed since the start of the session, we organized the data for each mouse into a 3D tensor. The tensor entries specified the number of video frames in which a particular syllable was performed at a specific location within a certain time epoch. Then, we developed a custom, nonnegative Tucker decomposition^42,43^ to approximate each mouse’s behavioral data as a weighted combination of shared factors (**Methods**). Like in a singular value decomposition of a matrix, each shared factor was a low-rank combination of *syllable*, *position*, and *epoch factors*, which specified what, where, and when syllables were used, respectively (**Fig. 2a**). Finally, the *mouse weights* specified how to combine these shared factors to best approximate each mouse’s behavioral tensor. We used cross-validation to determine that 22-dimensional mouse weights, 22 syllable factors, 4 position factors, and 4 epoch factors best explain the data (Extended Data Fig. 3, Extended Data Fig. 4a–c).

The epoch, position, and syllable factors captured low-dimensional patterns of syllable usage (**Fig. 2b**). For example, the first epoch factor showed an exponential decay over the first 20 min of the session, which captured transient behavior after mice were initially placed in the arena (**Fig. 2b**, top left, red line). The second epoch factor ramped up as the mice became accustomed to the environment; the third captured the sudden change when the lights turned on; and the fourth captured the last 15 min when the lights were turned off again (**Fig. 2b**, top left). The position factors tiled locations near the center, middle, and periphery of the arena, allowing the model to capture different syllable usage at different locations (**Fig. 2b**, bottom left). The syllable factors captured sets of syllables that tended to be used in similar locations and time epochs, like subsets of pausing syllables (**Fig. 2b**, right).

The mouse weights offered a condensed summary of each mouse’s syllable usage across space and time (Extended Data Fig. 4d). The weights showed considerable variability across drug groups, with 2AC+, 2A–, 2C–, 1A+, and 1B+ placing high weights on distinct subsets of shared factors (**Fig. 2c**). The other drug groups (2A+, 2C+, 1A–, and 1B–) were more similar in their weights to the saline control weights (**Fig. 2c**). To quantify how much these behavioral summaries differ across drug groups, we trained a multiclass logistic regression to predict which drug a mouse was given. The classifier achieved 45.8% accuracy on held-out mice, compared to a baseline of 10% accuracy from random guessing. The confusion matrix showed that the 2AC+, 2A–, 2C–, 1A+, and 1B+ groups were indeed much easier to distinguish than others (**Fig. 2d**). Differences in overall syllable usage were most informative, but the best classifiers also leveraged spatiotemporal information, as encoded by the weights of the behavioral model (Extended Data Fig. 4e).

The tensor decomposition also shed light on how drug groups differ in their patterns of syllable usage. The three most informative factors for drug discrimination best described the 1B+, 1A+, and 2AC+ drug groups, respectively (**Fig. 2e**). The first factor was mostly used by mice in the 1B+ treatment group, and it emphasized locomotion syllables near the edge of the arena that decreased in usage over the four major epochs—a finding that corroborates the scalar analyses (**Fig. 1**). The second factor, utilized mostly by 1A+ and 2A– mice, indicated a shift from pausing behavior to a single slow walking syllable over the course of the behavioral session. The third factor was predominantly featured in 2A+, 2AC+, and 2C– treated individuals and comprised some of the same locomotory syllables as in factor 1, but also a set of epoch-1-associated, wall-oriented rearing behavior and locomotion in the center of the arena. Combining the shared factors with the weights of the multiclass logistic regression showed which aspects of behavior were most predictive of each drug group (Extended Data Fig. 4f). This behavioral model decomposes spontaneous mouse behavior into shared factors, each encoding a distribution over what, when, and where syllables are used. Each mouse uses a unique mixture of factors, as encoded in their weights. The weights not only summarize spontaneous behavior and highlight differences among treatment groups, but also facilitate comparisons between behavior and the Fos maps described below.

### Effects of drugs on whole-brain Fos intensity

To determine the effect of serotonin receptor agonists and antagonists on neuronal activity, we used whole-brain Fos immunoreactivity as an approximate measure of integrated activity over the period when the mice were exploring the open field under each drug’s influence (Fos proteins usually peak ∼1–2 hrs after neuronal activation^44^). While sacrificing the temporal dynamics of neuronal activity recording, this approach provides spatial resolution of individual neurons at the scale of nearly the entire brain. Our pipeline (**Fig. 3a–c**) identified on average 7.3 million nuclei (**Fig. 3d**) with continuous Fos intensity levels in addition to binary ON/OFF counts used by most studies. Fos counts and intensity are positively correlated, but with considerable variation along both axes (**Fig. 3e; Methods**). The Fos maps were all registered to the Allen Brain Atlas^45^, allowing averaging within brain regions. **Fig. 3f** shows a representative coronal section of Fos intensity above (red) or below (blue) those in the saline control (presented as raw Fos intensity map in **Fig. 3f_10_**) for 9 drug groups (**Fig. 3f_1–9_**). Each Fos map is an average of ∼17 mice, with individual mice exhibiting different degrees of variation in different drug/brain region combinations (**Fig. 3g, h; Extended Data** Fig. 5). The effect of the drugs on the Fos intensity along entire anterior-posterior axis of the brain can be viewed in **Movie S1**.

In principle, the patterns of Fos map modulation by agonists and antagonists should reflect the expression patterns of the cognate receptors. However, the following factors make such reflection less straightforward. First, in addition to directly acting on receptor-expressing cells, drugs could also have indirect effects on cells that do not express the receptor. Second, some serotonin receptors such as 1A and 1B are known to be enriched on axon terminals^25,26^, such that even direct effects may be exerted on brain regions distinct from receptor-expressing cells. Third, agonists can only have an effect when receptors are not fully activated by the behavior, while antagonists can only have an effect when receptors are activated by the behavior. With these caveats in mind, we use **Fig. 3f** to illustrate a comparison between agonists and antagonists against the same receptor (see **Table S2** for potential explanations of different patterns). (1) The agonist and the antagonist modulated Fos intensity in opposite directions, as is the case for the 2A+ and 2A– in the somatosensory cortex (**Fig. 3f_1_** and **3f_2_**, arrows). (2) Only the agonist substantially modulated Fos intensity, as is the case for the 1A+ and 1B+ in bed nucleus of stria terminalis (BNST; **Fig. 3f_5_** and **3f_7_**, arrowheads) or the 1B+ in dorsal striatum (**Fig. 3f_7_**, arrow). (3) Only the antagonist substantially modulated Fos intensity, as is the case for 2A and 2C antagonists in BNST (**Fig. 3f_2_** and **3f_4_**, arrowheads). (4) Both the agonist and antagonist modulated Fos intensity in the same direction, as is the case for 2C in somatosensory cortex (**Fig. 3f_3_** and **3f_4_**, arrows). (5) Neither the agonist nor the antagonist modulated Fos intensity, as are the cases for the 1A and 2C agonists and antagonists in the dorsal striatum.

**Fig. 4.**
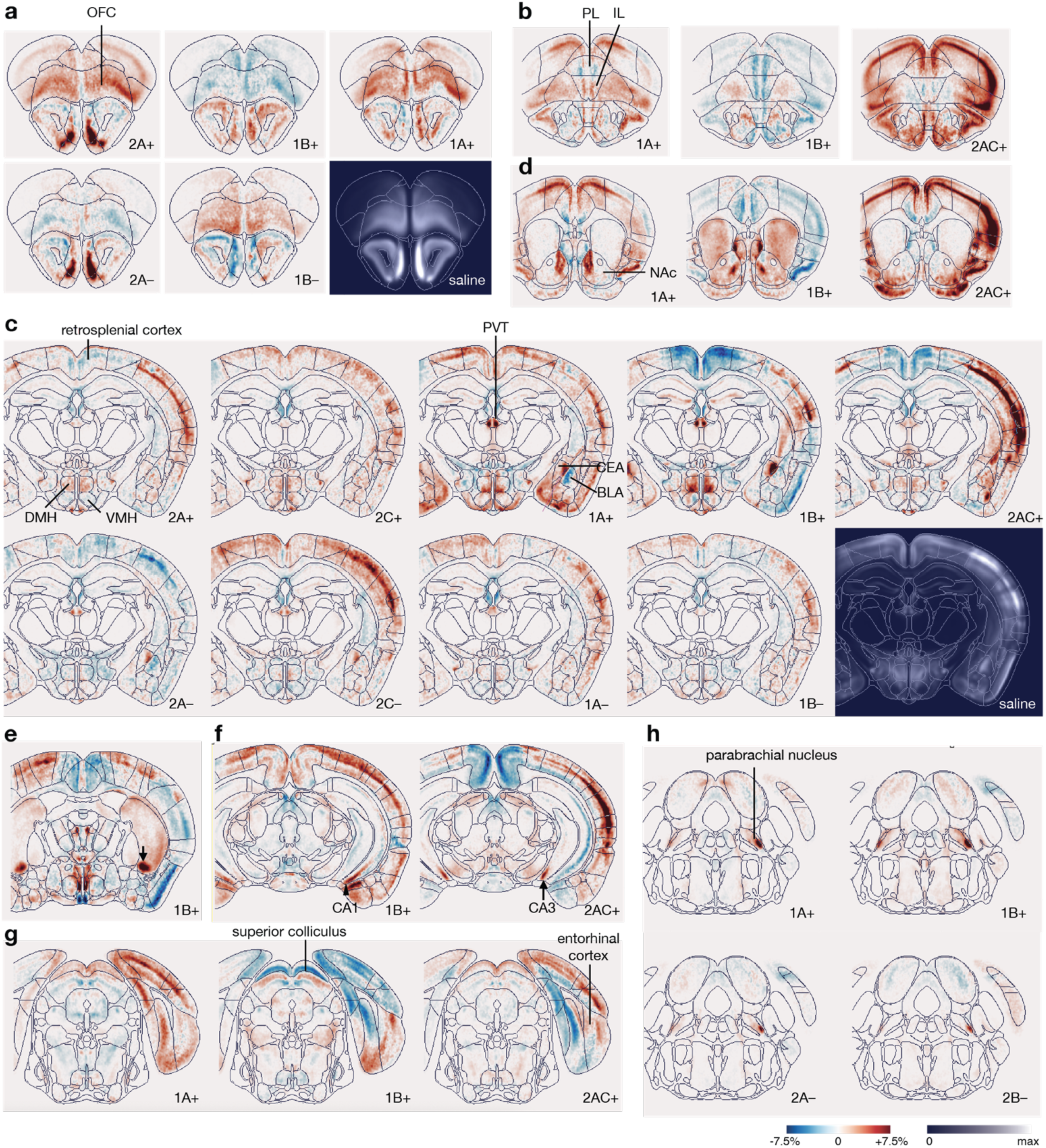
Examples of the effects of serotonin receptor agonists and antagonists on Fos intensity in selected regions. All images are saline-subtracted Fos intensity maps from 100-µm coronal slices highlighting specific brain regions that are modulated by specific subsets of agonists and antagonists as indicated at the bottom right. Heat maps same as Fig. 3f. **a,** Orbitofrontal cortex (OFC). **b,** Medial prefrontal cortex, including prelimbic (PL) and infralimbic (IL) areas. **c,** A central section highlighting drug actions on the retrosplenial cortex, dorsomedial and ventromedial nuclei of the hypothalamus (DMH and VMH), paraventricular thalamus (PVT), and central and basolateral amygdala (CEA and BLA). **d,** Nucleus accumbens (NAc). **e,** Ventral tip of posterior striatum (arrow). **f,** Ventral tip of hippocampal CA1 (arrowhead) and CA3 (arrow) areas. **g,** Superior colliculus and entorhinal cortex. **h,** Parabrachial nucleus. See text for detailed description of modulation by specific agonists and antagonists.

In some cases, our Fos maps are consistent with direct modulation of receptor-expressing cells based on expression patterns of serotonin receptors^24,26–28^. For example, 2A is broadly expressed in excitatory neurons of the neocortex^28,46^, consistent with the strongest effect of 2A agonist in cortex. Among the four serotonin receptors examined here, 1B is most prominently expressed in dorsal striatum^24^, accounting for the strongest effect of 1B agonist on Fos intensity. Given that 1B is expressed predominantly in GABAergic spiny projection neurons in the striatum^28^, and activation of 1B hyperpolarizes receptor-expressing cells, one likely possibility for the 1B+ effect is that it is mainly exerted at presynaptic terminals of spiny projection neuron collaterals within the striatum, thus reducing GABA release. Indeed, experimental data from a recent report support this interpretation^47^. **Fig. 3g** offers another view of spatial regulation of Fos intensity across the brain by two of the nine drugs, 2AC+ and 1B+, with quantifications across all mice in 282 Allen regions (**Fig. 3h**; see Extended Data Fig. 5 for similar data for the rest of the drugs). The rich patterns of Fos intensity across the entire brain and drug conditions (**Fig. 3f–h, Extended Data** Fig. 5**, Movie S1**) provide a resource for exploring the effect of activating or inhibiting specific serotonin receptors on brain activity. We highlight a small subset of specific findings from our Fos maps in the next section, with a bias towards brain regions in which Fos intensity was modulated by multiple drugs and for which atlas-level summaries inadequately capture the spatial specificity of the modulation.

### Effects of drugs on Fos intensity in select brain regions

The prefrontal cortex, including the orbitofrontal cortex (OFC) and medial prefrontal cortex (mPFC), is strongly innervated by raphe serotonin neurons and in turn provides strong monosynaptic input to raphe serotonin neurons^48–51^. We found that Fos intensity in OFC was up-regulated by 2A+, 1A+, 1B–, but down-regulated by 2A– and 1B+ (**Fig. 4a**). mPFC is traditionally divided into prelimbic and infralimbic cortex (PL and IL). Interestingly, while 1B+ decreased Fos intensity in both regions, 1A+ decreased Fos intensity in PL but increased Fos intensity in IL, whereas 2AC+ increased Fos intensity in PL but decreased Fos intensity in IL (**Fig. 4b**). Thus, serotonin appears to modulate prefrontal cortex activity through multiple receptors with exquisite regional specificity in magnitude and direction.

Exquisite modulation by multiple serotonin receptors was also evident in many other brain regions. For example, while 2A+ generally increased Fos intensity across the neocortex, it decreased Fos intensity in the retrosplenial cortex. This effect was more pronounced with 2AC+ (**Fig. 4c**). Likewise, in the hypothalamus, while 2C+ increased Fos intensity in both dorsomedial and ventromedial hypothalamic nucleus (DMH and VMH), 2A+ increased Fos intensity in DMH but decreased Fos intensity in VMH, and 1A+ and 1B+ modulated Fos intensity in specific subregions of DMH and VMH (**Fig. 4c**). In the thalamus, 1A+ and 1B+, along with 2A– and 2C–, all increased Fos intensity in a bilaterally symmetric pair of hotspots in a specific part of the paraventricular thalamic nucleus (PVT) (**Fig. 4c**). In the amygdala, 1A+ and 2A– increased Fos intensity in the lateral portion of the central amygdala (CEA) while decreased Fos intensity in the adjacent medial portion of the basolateral amygdala (BLA), whereas 2C– antagonist and 1B+ exhibited a similar effect only on lateral CEA but not on medial BLA (**Fig. 4c**). The above examples suggest heterogeneity exists within a given brain region in the Allen Brain Atlas as revealed by Fos intensity modulation by serotonin receptor agonists and antagonists. Such sub-region specificity was found in many other brain regions. For example, 1A+, 1B+, and 2AC+ modulated Fos intensity in distinct subregions of the nucleus accumbens (**Fig. 4d**). 1B+ increased Fos intensity highly in the ventral tip of posterior striatum (**Fig. 4e**, arrow). In the ventral hippocampus, 2AC+ increased Fos intensity preferentially in the ventral tip of CA3 (**Fig. 4f**, arrow), whereas 1A+ increased Fos intensity preferentially in the ventral tip of CA1 (**Fig. 4f**, arrowhead). Furthermore, 1A+, 1B+, and 2AC+ modulated Fos intensity in the superior colliculus in a layer-specific manner and in the entorhinal cortex in a subregion-specific manner (**Fig. 4g**). Finally, 1A+ and 2A+, as well as 2A– and 2C–, increased Fos intensity in the ventrolateral subregion of the parabrachial nucleus (**Fig. 4h**), which relay body information to the brain. To our knowledge, specific functions of some of the above subregions have not been described. Examination of serotonin receptor expression from spatial transcriptomic dataset^27,28^ did not reveal obvious subregion-specific expression that correspond to subregion-specific Fos intensity maps. These findings illustrate the effect of serotonin modulation of highly specific brain regions, consistent with the highly specific collateralization patterns of subpopulations of serotonin neurons^51,52^, and suggest that differential serotonin modulation can be used as a means to subdivide brain regions.

Overall, the effects of 2A+ and 2A– were often in opposite directions, with the 2A+ increasing and 2A– decreasing Fos intensity. This is consistent with that fact that 2A is coupled to G_q_ and thus its activation should depolarize target neurons directly innervated by serotoninergic axons. 1A+ and 1B+ gave some of the largest magnitude of modulation, often with remarkable region specificity (**Fig. 3f**, **Fig. 4, Extended Data** Fig. 5). While 1A and 1B are both coupled to G_i_ and thus their activation should hyperpolarize direct target neurons, 1A+ and 1B+ often increase Fos intensity in some brain regions. This could be because they may act on local inhibitory neurons or axons of long-distance projecting inhibitory neurons innervating the target regions. Curiously, 1A+/1B+ often modulated Fos intensity similarly as 2A–/2C– (e.g.: BNST, **Fig. 3e**; lateral CEA, **Fig. 4c**; parabrachial nucleus; **Fig. 4h**); sometimes 2A– more closely mimicked 1A+ and 2C– mimicked 1B+ (e.g.: medial BLA, **Fig. 4c**), whereas other times it was the other way around (e.g.: OFC, **Fig. 4a**; retrosplenial cortex, **Fig. 4c**). These observations suggest cooperation between different combinations of serotonin receptors. We note that the similarities of 1A+ and 2A– effects were also observed in behavioral analyses (**Fig. 1g–i**).

### Decomposition of Fos maps

Fos imaging yields high-dimensional, whole-brain maps for each mouse. We hypothesized that the application of serotonin receptor agonists and antagonists would produce correlated changes in neural activity, which would manifest in low-dimensional, spatial structure in the whole-brain Fos maps. To test this hypothesis, we constructed a model tailored to Fos measurements that can find low-dimensional factors of variation across the collection of mice (**Fig. 5a**). Conceptually, we think of the Fos measurements as a marked spatial point process: each cell has a location (3D) and an intensity (positive scalar). The point process intensity function specifies both the expected number of Fos cells and the expected Fos intensity for each location in the brain. For computational tractability, we approximated the intensity function as piecewise constant within (100 µm)^3^-voxels (and each brain comprises 350k such voxels). We modeled the intensity function as a weighted combination of nonnegative factors. Each factor is normalized, so that it can be thought of as a distribution over voxels. For each mouse, we jointly modeled the expected Fos cell count and intensity as weighted combinations of these factors (**Fig. 5b, c, Extended Data** Fig. 6a**; Methods**).

**Fig. 5.**
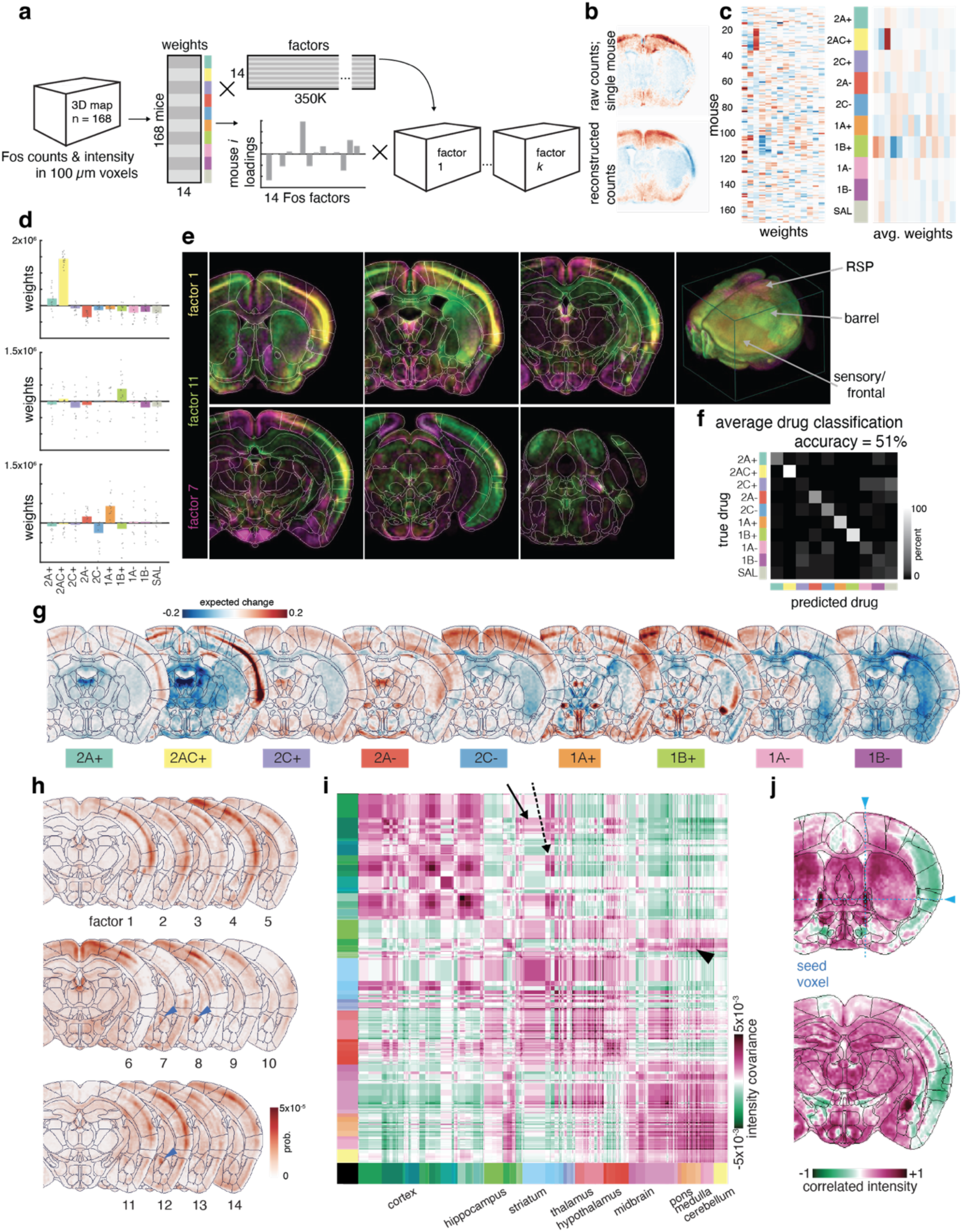
Decomposition of Fos maps. **a**, Schematic representation of the semi-non-negative matrix factorization of Fos intensity and count data. All raw data brain maps are factorized into per-mouse weights and 14 separate non-negative Fos factor maps. **b,** Example coronal slice from a saline-subtracted Fos map for a single mouse and the model reconstruction. **c,** The Fos count weights for each mouse (left) and the average for each drug group (right). **d,** The weights for three example factors show differential usage across drug groups. **e,** 2D (coronal slices at different anterior to posterior positions, left 6 panels) and 3D (right panel) visualizations of these three factors highlight regions of overlap and complementarity. RSP, retrosplenial cortex. **f,** Confusion matrix of drug classification based on weights from the Fos model. The average accuracy is 51%, well above the baseline prediction of 10% given 10 groups. **g,** Weights from the logistic regression combined with the factors of the Fos model show which changes in Fos activity relative to saline are most predictive of each drug. **h,** Single coronal slice from all 14 factors. Arrowhead indicates a region of interest in central amygdala which is apparent in 3 separate factors. **i,** Brain-wide covariance map (as estimated by the 14-dimensional factor model) summarized at the level of Allen Brain Atlas regions. Regions are scaled proportionally to their volume and entries on the diagonal represent the mean covariance of the voxels within that region. Solid arrow, motor cortex and striatum; dashed arrow, frontal cortex and amygdala; arrowhead, subiculum and pons. **j,** A single row of the brain-wide correlation matrix from a seed voxel in BNST highlights the regions and networks that may be co-regulated with BNST, including CEA. Voxel values are correlations from –1 to +1.

As with the behavioral syllable usage, we found that factor weights describing Fos counts differ substantially between drug groups. For example, weights on factor 1 are largest among mice given 2AC+ (**Fig. 5d**), and factor 1 places highest probability on voxels in the frontal cortical regions (**Fig. 5e**). Factor 11 places higher probability on the striatum and barrel cortex and is most used by mice given 1B+. Factor 7 is most used by mice given 1A+ and places highest probability on the retrosplenial cortex (RSP) and ventral premammillary nucleus (**Fig. 5d, e**).

Factor usage is a strong indicator of drug group, and given the count and intensity weights, we can predict which drug a mouse was given with 51% accuracy using a multiclass logistic regression (**Fig. 5f**). The weights of the logistic regression offer insight into which voxels are most predictive of each drug. **Fig. 5g** shows the combined factors obtained with the logistic regression weights best predicting each drug group. For example, in the depicted coronal section, increased activity in cortex and decreased midline thalamic activity is most predictive of drug 2AC+, whereas increased activity in caudoventral striatum is most predictive of drug 1B+.

Interestingly, our model reveals hot spots of correlated activity at finer resolution than the Allen atlas affords. By emphasizing the changes filtered by the drug classification task, voxels that are informative for predicting treatment group are highlighted even when the magnitude of their mean saline subtracted intensity is small. In many such cases, the model expects changes in opposite directions for agonist/antagonist pairs even though information on direction of effect was not included in the regression. For example, in the pontine central gray, parabrachial nucleus was previously highlighted in **Fig. 3** and **Fig. 4**; however, the factorization also expects changes in Barrington’s nucleus (opposing direction of change in 1A+ vs. 1A–) and nucleus incertus (opposing direction of change in 1A+ vs 1A–, and 1B+ vs. 1B–; Extended Data Fig. 6b). Amygdalar hot spots in the 2AC+ treated animals are similar to those in the 2A+ condition; a sign inversion in the modeled changes of 2A– treated animals matches in these spots (Extended Data Fig. 6c).

Since some of these subregions appear in only a subset of the factors (e.g., CEA, **Fig. 5h**), the maps of specific factors could be considered as being ‘coactive’ with that subregion. To further dissect this possibility, we generated a model-based approximation of the voxel-by-voxel covariance matrix across the entire brain. However, it is infeasible to visualize the entire 350k x 350k covariance matrix that we obtain with (100 µm)^3^-voxels. Instead, to present a comprehensive summary of brain-wide covariance, we first computed the average covariance for all pairs of Allen atlas regions to obtain the intensity-based covariance matrix in **Fig. 5i** and counts-based covariance matrix (Extended Data Fig. 7). Values on the diagonal depict covariance of intra-regional voxels. At this resolution, three large clusters of correlated regions emerged: one cluster consisting of cortical regions; a second cluster of hippocampus, striatum, thalamus, and hypothalamus; and a final cluster of midbrain, pons, medulla, and cerebellum. While some of these relationships are expected (e.g., motor and frontal cortices preferentially correlating with striatum and amygdalar regions, respectively), long-range and less-tested correlations will require experimentation to validate (e.g., pons with subiculum).

Finally, to visualize fine-grained correlation structure at the level of individual voxels, we normalized by the standard deviation to obtain a 350k × 350k correlation matrix, and then examined individual rows of that matrix. Each row can be reshaped into a 3D map that illustrates the correlation between all voxels and a single “seed” voxel (**Fig. 5j**). Selecting a single source in dorsal BNST, we see that this voxel is highly correlated with its neighbors, as expected from the hot spot in the corresponding factors. Less apparent is that this voxel is highly correlated with activity in the dorsal striatum, PVT, and a hotspot in the central amygdala, and it is anticorrelated with activity in somatosensory cortex. Given that the high correlation between BNST and central amygdala is consistent with the known functional similarity between these regions^53^, our finding suggests an approach for hypothesizing relationships between less well-studied regions. We have developed an interactive viewer to explore brain-wide, (100 µm)^3^-resolution correlations like these, and several examples are shown in Extended Data Fig. 6d.

### Joint neuro-behavioral modeling and prediction

Thus far, we have described the effects of activating or inhibiting specific serotonin receptors on animal behavior (**Fig. 1** and **Fig. 2**) and separately on integrated neuronal activity as measured by whole-brain Fos intensity (**Fig. 3–5**). Interestingly, both types of measurements yielded similar accuracy when predicting which drug an animal was given (**Fig. 2d** and **5f**). To better understand the relationship between behavior and brain activity, we next developed joint models to predict one from the other, leveraging paired behavioral and Fos data for each of the 168 animals.

First, we looked for correlations between the per-mouse weights from our behavioral factor model (**Fig. 2**) and our Fos factor model (**Fig. 5**). We used canonical correlation analysis (CCA) to identify pairs of directions in behavioral and neural weight space that exhibited the strongest correlation. We found that the top three canonical directions had correlation coefficients of 0.82, 0.70, and 0.66, respectively (Extended Data Fig. 8b), suggesting a strong relationship between the weights of our two models. The CCA directions captured relationships between neural activity and behavior: for example, the first pair of canonical directions captured a correlation between rearing near the periphery of the arena and activity in motor cortex (Extended Data Fig. 8a).

Can we predict time-integrated neural activity from behavior, or vice versa? To do so, we fit a linear model to predict the weights of our behavior factor model given the weights of the Fos factor model (**Fig. 6a**). We trained the linear model on the weights from 75% of the mice and evaluated the predictions on the remaining 25% (**Methods**). With the Fos weights, the linear model can explain a greater fraction of deviance than our baseline behavioral model. The degree of improvement varies across drug groups, with the 2AC+, 2A–, 1A+, and 1B+ groups showing the greatest increases (**Fig. 6b**). We compared against several baseline models to show that the predictive power in the Fos signal is primarily derived from differences in both Fos and behavior across drug groups (**Methods, Extended Data** Fig. 8c). We also trained models to predict Fos weights given behavior weights and found that predictions were less accurate in this direction. Only the 2AC+ and 1B+ groups showed improvements over baseline when given behavior information (Extended Data Fig. 8d). The poor predictive performance for other drug groups suggests that Fos weights contain additional information about variation in neural activity that is not reflected in overt behavior, at least as measured by syllable usage across time and space. Next, we used the linear models to predict responses to synthetic “perturbations” of behavior or neural activity (**Methods**). Given the limited performance of the linear mapping from behavior to Fos weights, care must be taken when interpreting predictions in this direction. Nevertheless, our model predicted increased pausing to correspond with a decreased Fos activity (i.e., fewer Fos positive cells) in the striatum (Extended Data Fig. 8e, f). Likewise, we could predict changes to the Fos map if we increase syllable frequencies for rearing, grooming, and locomotion (Extended Data Fig. 8g). Interestingly, an increase in grooming syllables predicted an increase of Fos activity in the somatosensory cortex, whereas an increase of locomotion syllables predicted an increased Fos activity in the striatum, opposite of that of the pausing syllables. This latter prediction is consistent with experimental data that both direct and indirect pathway spiny projection neurons, which comprise most of the striatal neurons, increase their activity upon locomotion onset^54,55^.

**Fig. 6.**
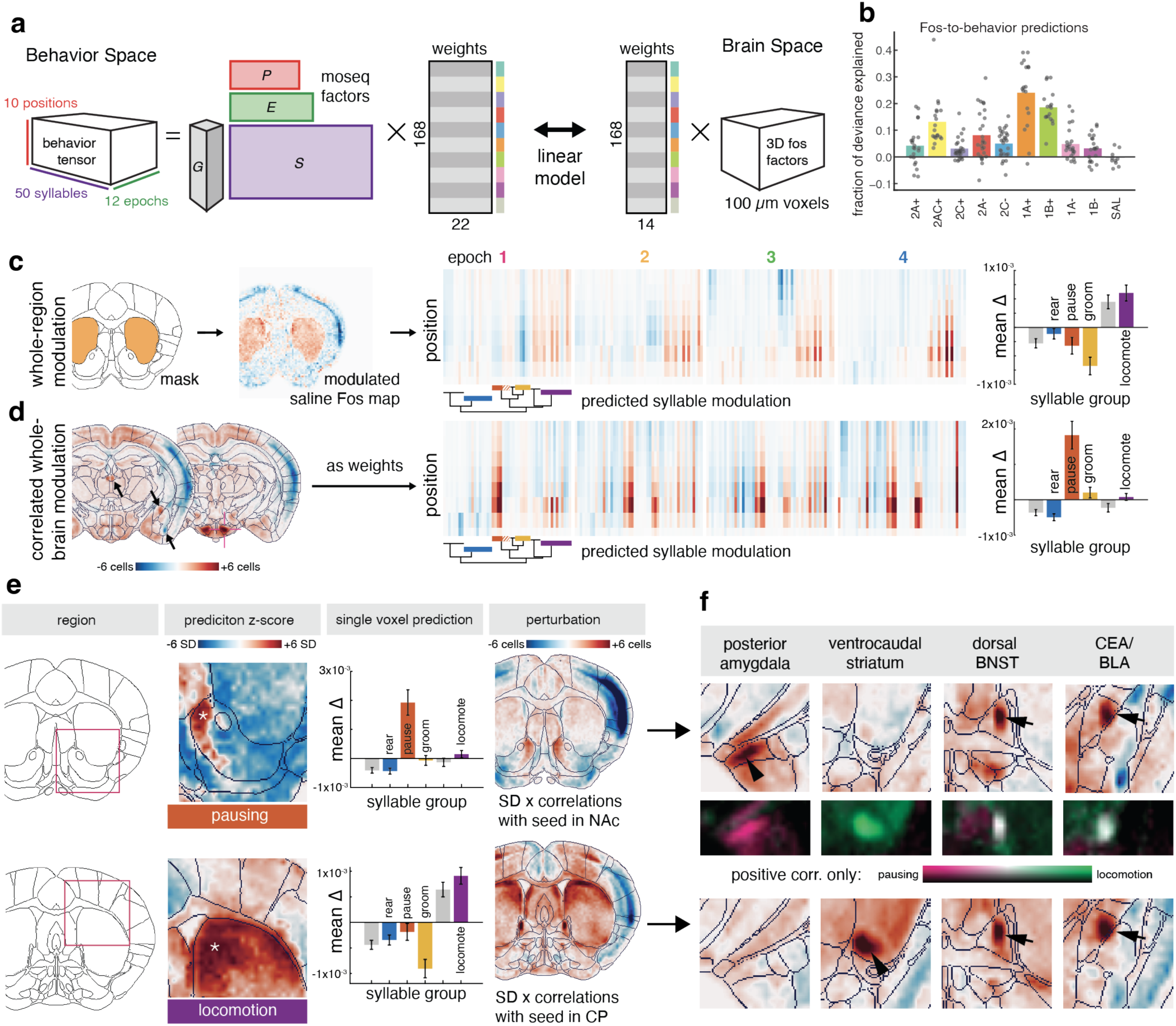
Linking behavior and Fos map. **a**, Schematic of the strategy to link behavior and Fos map using linear models. **b,** Predictive power of Fos-to-behavior linear model depicted as fraction of deviance explained from held out test mice, relative to a baseline model that predicts the average syllable counts (Methods). **c,** Predicting behavior from synthetically upscaling an average saline brain’s Fos activity in the striatum. Following the right-to-left path in **a** yields predictions of Fos factor weights to estimate the manipulated brain, predictions of the behavior factor weights from the regression, and predictions of the corresponding behavioral tensor, here shown as syllables × position matrix across the 4 major epochs. Bar plot (right) shows the total change in major behavior groups from the unmodulated saline brain’s syllable predictions with 95% bootstrap confidence intervals. **d,** Correlated whole-brain modulation changes cell counts across the entire saline average brain in proportion to the correlation with a seed voxel (see **Fig. 5j**). Here, a seed voxel in ventral premammillary leads to strong modulation in correlated regions like PVT, CEA, and ventral BLA, as indicated by arrows. This pattern of modulation predicts increased pausing across all epochs. Bar plot (right) similar to (**c**). **e,** With a focus on the nucleus accumbens (NAc) (top row) and dorsomedial striatum (DMS) (bottom row) voxel-wise z-scores for predictions of individual behavioral categories reveal a significant hotspot for predicting pausing behavior in NAc and a significant portion of DMS as predicting locomotion. Selecting a single seed voxel in each hotspot and repeating the process in (**d**) highlights the selectivity of each region in predictions across all behaviors as well as the correlated and anti-correlated brain regions included in the perturbation. **f**, Four highlighted hotspots in maps of correlation with the seed voxels in (**e**) reveal posterior amygdala as positively correlating with NAc, ventrocaudal striatum as correlating with DMS, while dorsal BNST and CEA positively correlate with both NAc and DMS in spite of their differential behavioral predictions.

Finally, we simulated the effect of synthetic perturbations of Fos activity on behavior, the direction in which our linear models showed greater predictive performance. Increased Fos activity in the striatum (**Fig. 6c**, left) predicted a broad increase in locomotion toward the periphery of the arena, at the expense of a decrease in rearing, pausing, and grooming (**Fig. 6c**, middle and right; **Methods**), consistent with results from our behavior-to-Fos-map prediction (Extended Data Fig. 8e, f). We performed a stratified bootstrap analysis to obtain confidence intervals for these predictions (**Methods**). As Fos is a static time-integrated signal, predicting epochs for individual behaviors is challenging to interpret. However, it is important to note that the behavioral protocol was the same for all sessions; in particular, the LED turned on and off at the same time. Specific patterns of Fos may correlate with stronger responses to the LED or a greater tendency to habituate in the arena, both of which would appear in the distribution of syllables across epochs. The striatal perturbation in **Fig. 6c** predicts increasing locomotion relative to saline across epochs, which when considering 1B agonism as the leading driver of this pattern, follows from this group’s reduced habituation of speed relative to saline (**Fig. 1c**).

We also considered “correlated whole-brain modulation” to mimic a perturbation of a functional network of brain regions. Starting with a seed voxel, we modulated all other voxels up or down depending on their correlation with the seed voxel and the brain-wide variability in Fos (as in **Fig. 5j**). Each voxel’s perturbations are of up to ±1 standard deviation from the average saline mouse’s value at that voxel (**Methods**). As an example, choosing a voxel in ventral premammillary (PMv) led to a brain-wide modulation that included increased activity in central amygdala (CEA) and decreased activity in cortex (**Fig. 6d**, left). This synthetic perturbation was predicted to correspond to an increase in pausing behavior (**Fig. 6d**, right).

We repeated the analysis for all 350k voxels throughout the brain to provide a resource of behavioral predictions starting from any seed voxel (Extended Data Fig. 9a). For example, seed voxels in dorsal striatum predicted the largest increases in locomotion (**Fig. 6e, Extended Data** Fig. 9b), consistent with findings of complementary activity in the direct and indirect pathways during movement^54,55^, and the largest predicted increases in pausing localize to seeds in anterior medial NAc (**Fig. 6e**). Having identified the seeds for predicting high magnitude changes in each major behavioral category (Extended Data Fig. 9b), we then identified correlated hotspots in other regions that contribute to these predictions. One hypothesis would suggest that pausing and locomotion should be predicted by entirely separate networks; however, the absence of a dynamic signal in Fos imaging does not preclude the possibility of overlap in these correlated regions. Indeed, posterior amygdala and ventrocaudal striatum specifically correlate with pausing and locomotion networks respectively, but BNST and CEA each correlate with both networks (**Fig. 6f**).

Importantly, this model makes predictions about the full behavioral tensor for each syllable-position-epoch combination, which can be averaged across categories of interest as above (Extended Data Fig. 9c). Additionally, individual syllables in a specific set of positions and epochs can be interrogated from each seed voxel in the brain. Across bootstrapped model fits, many seed voxels have varied or inconsistent predictions for individual syllables, but many rearing syllables are well-predicted to increase with perturbations originating from seed voxels in cortex. Three of four pausing syllables are well-predicted by seeds in lateral septum, and individual locomotory syllables are differentially predicted by seeds in striatum and cortex (Extended Data Fig. 9d).

Currently, brain-wide perturbations like these are beyond our experimental capability, but as our genetic toolkit for sophisticated perturbations of neural activity continues to grow, our joint neuro-behavioral modeling offers a means of generating hypotheses for how perturbations of one modality will manifest in the other.

## DISCUSSION

Here we report the effect of agonists and antagonists of four widely expressed serotonin receptors on spontaneous mouse behavior as well as on Fos intensity as a proxy of integrated activity at the scale of the entire mouse brain. We used custom factor analyses, which were tailored to the high-dimensional Fos and behavioral measurements. These models yielded lower dimensional representations of the complex data that shed light on the dominant factors of variation in each domain, capture predictable differences across drug groups, and enabled predictions of behavioral changes following perturbations of the Fos maps and vice versa.

Agonists and antagonists of specific serotonin receptors have been extensively tested in rodents, usually on behaviors aimed at modeling human psychiatric disorders such as anxiety and depression, as selective serotonin reuptake inhibitors are the most widely prescribed medications for their treatment^56,57^. To our knowledge, none of the drugs we used have been examined on spontaneous behaviors using modern analysis methods such as MoSeq, which was demonstrated to outperform traditional scalar measures to distinguish the effect of different drugs^41^. Furthermore, with the exception of 2AC+, we chose our agonists and antagonists to be minimally cross-interfering with non-intended receptors^33–40^. Thus, our parallel studies of 9 different drugs using both scalar and MoSeq analyses provide a valuable resource on the effect on animal behavior in response to preferentially activating or inhibiting four widely expressed serotonin receptors.

Recent technological advances in imaging and whole-mount histology have enabled examination of Fos (or products of other immediate early genes) at the scale of the entire mouse brain^58–63^. We integrated these methods with deep learning–based image extraction^64^, enabling us to identify not only an average of 7 million Fos+ cells per brain but also determine their intensity levels. The comprehensiveness of whole-brain Fos map changes in response to specific serotonin receptor agonists and antagonists enabled many new biological discoveries (**Fig. 3** and **Fig. 4**) and provided a rich resource for the field—all data are freely available to the research community. We note that the effects of activating or inhibiting serotonin receptors on Fos maps we report here correspond to integrated activity of mice performing spontaneous behavior in an open arena, including a period with LED light on for heightening anxiety. The effects of manipulating serotonin receptor activity (or other manipulations) on Fos maps corresponding to mice performing other behaviors can be similarly obtained using our data acquisition and analysis pipelines (**Fig. 3**). It will be of great interest in the future to compare the effects of manipulating serotonin receptor activity in different behavioral paradigms.

Our Fos map analyses repeatedly identified subregions of serotonin modulation within the regions defined by Allen Brain Atlas. To our knowledge, many of these subregions (e.g., the ventroposterior corner of the striatum activated by 1B+, **Fig. 4e**) have not been described in the literature for their unique anatomical, physiological, or functional properties. We did not find correlation of such “modulation hotspots” with receptor expression from high-resolution spatial transcriptome datasets^27,28^. Other possible explanations for such modulation hotspots could be due to differences of these subregions compared to their neighbors in: 1) density of serotonin fibers; 2) expression levels of cognate serotonin receptors in presynaptic terminals of long-distance axons; 3) differential engagement in some behavior mice perform in our assay; 4) combination of the above and/or indirect effects. Regardless of the detailed mechanisms, these modulation hotspots highlight that studying serotonin neuromodulation can help dissect functional heterogeneity of diverse brain regions.

Our factor analyses of behavior using tensor decomposition enabled us to not only include MoSeq syllables, but also the position and the temporal epoch in which these syllables were performed. Likewise, our factor analysis of Fos map using a marked spatial point process model enabled us to predict drugs based on Fos maps and which brain regions are most predictive of specific drugs. Furthermore, the factor model yielded an estimate of the brain-wide covariance map, which captures how changes in Fos intensity in one voxel correlates with changes throughout the rest of the brain. The diversity of serotonin drugs helped drive a range of neural activity patterns to probe this covariance structure. While these maps are specific to this dataset, adding new samples or external datasets could further expand upon or refine the patterns already present. Among other utility, this covariance map aids in prediction of behavioral changes due to changes in Fos maps, as we discuss next.

Low-rank factor analyses of spontaneous behavior and Fos maps enabled us to explore their relationship. Neural encoding and decoding models have shown impressive abilities to relate real time behavior and neural activity^65–67^, but it is unclear *a priori* whether behavior measured in real time with temporal resolution of several hundred milliseconds (the duration of average MoSeq syllables) can be related to the Fos map, corresponding to a single snapshot of integrated activity over tens of minutes. Canonical correlation analysis and predictive modeling revealed a strong correlation between the two modalities, demonstrating rich information contained in the static Fos maps. Importantly, when considering both the low dimensional factorizations and the cross-modal predictions, none of the models included drug identity information or optimized for classification performance. Drug prediction accuracy following behavioral decomposition nearly matches the full dataset’s predictive power, with just 14 Fos factors predicting these 9 drugs, all serotonergic, and some with overlapping targets, well above chance. Importantly, the use of 9 different drugs led to greater variation in behavior and Fos activity than occurs naturally in the saline control group, and the increased dynamic range should improve estimates of neural-behavioral relationships. Some of these predictions are consistent with previous experimental data (e.g., increased striatal activity leads to increased usage of locomotion syllables and vice versa) or intuition (e.g., increased grooming leads to increased somatosensory cortex activity).

The mutual predictions between behavioral syllable usage and Fos maps enable researchers to explore the relationship between brain and behavior and generate specific hypothesis for future experimental validation. While many brain regions of known function are placed into larger context in this joint dataset, we are especially interested in the hypotheses that stem from small subregions, including those that do not obey reference atlas borders and those that strongly correlate with each other across long distances. The whole-brain projections of serotonergic neurons and the diverse set of receptors provide a useful system in revealing surprising heterogeneity in downstream target regions and their interactions. Finally, we note that our cross-modality neuro-behavioral modeling strategy could be applied to analyzing relationship between other datasets with different time scales, such as behavior and gene expression profiling (e.g., from the single-cell/nucleus RNA-sequencing), as well as neural activity (real time or integrated Fos map) and gene expression.

## METHODS

### Mice

All animal procedures followed animal care guidelines approved by Stanford University’s Administrative Panel on Laboratory Animal Care (APLAC). Wild-type mice of the C57Bl6/J strain were used for all experiments (n = 190 8-week-old male mice from the Jackson Laboratories). 20 mice per week arrived in our animal facility and were housed on a reverse 12-h light/12h-dark cycle for 12–15 days. The afternoon before a cohort’s experiment, a cage of 5 individuals was split into 5 single-housed cages so that the last mouse of the group would not be in a more anxious state than others before behavioral testing.

### Drug preparation

Drug identities and concentrations are listed in **Table 1**. Briefly, stock solutions were made as recommended by the manufacturers in water, if possible. In cases where solubility was an issue, DMSO was used for the stock solution instead. Stock concentrations were diluted in saline (0.9% NaCl) immediately before loading into a syringe at an injection volume of 2 μl/g of mouse weight. The identity of the substance being injected was unknown to the behavioral experimenter. Before intraperitoneal (IP) injection, each mouse was very briefly sedated by isoflurane using a small volume of anesthetic on a nestlet inside an inverted chamber to minimize perceived handling and injection stress. Mice underwent an IP injection of one of the serotonin receptor agonists or antagonists, or saline, 30 min prior to behavioral recording. The drugs were counterbalanced so that each drug was present at each recording timepoint during the day-long session and over the days of the week of experimentation. The final drug solutions were always used on the day they were prepared.

### Behavioral assay

The camera and arena for behavior were constructed according to specifications listed on https://github.com/dattalab/moseq2-app/wiki. The overhead Kinect camera was frame synchronized via a simulated DAQ with two laterally mounted DMK 33UX273 monochrome cameras (The Imaging Source) and a central collimated LED (4900 K, 740 mW, #MNWHL4, #SM2F32-A ThorLabs, powered by a DC4104 driver, ThorLabs) and controlled by MicroManager software.

All behavioral experiments were performed by one female experimenter between the weekday hours of 10 am and 5 pm. The behavioral room was kept under red light and with white noise present. Each day 5 mice were tested, and no mouse was tested more than once. Each mouse was taken from the reversed light cycle room in a light-tight container and brought to the behavioral room 5 min before IP injection. Behavioral testing began 30 min after IP injection. After this post-injection wait period, the mouse was placed in the center of a round open arena (US Plastic, 14317). The mice were transferred to and from the arena via a 5-inch net pot to minimize handling stress and bedding transfer. In the arena, movement was unrestricted. Video recording was started immediately, and after 30 min, the LED was powered for 15 min, illuminating the center of the arena out to ∼65% the radius of the arena or ∼40% of the total area. After stimulus, the LED light was turned off and the mice had another 15 minutes of free movement throughout the arena. After cessation of recording, mice were transferred by cup back to their cage and left to rest in the same behavior room for 45 min to allow expression of Fos proteins before perfusion.

After each mouse, the arena was cleaned with 10% Alconox (VWR, 21835-032), water, and 70% ethanol, and left to fully dry before the next mouse was placed in it. One mouse was excluded from further analyses for failure to move in the arena and a second mouse was excluded due to a failure of the LED illumination.

### Behavioral analysis

Syllable extraction followed the MoSeq pipeline (https://github.com/dattalab/moseq2-app/wiki/MoSeq2-Extract-Modeling-Notebook-Instructions) (https://doi.org/10.1038/s41593-020-00706-3). Briefly, the 188 hours of recordings were processed to extract an oriented crop of the mouse from each full frame. Scalar information about the *x*–*y* position of the mouse, the height of the mouse, and the *x*–*y* area of the animal was also saved for each frame. The cropped data, 80 × 80 pixels × 108000 frames per mouse, were further reduced by principal components analysis (PCA), and subsequently all simultaneously fit by an auto-regressive hidden Markov model. Kappa, the free parameter that regulates the duration of individual states, was selected to fit a model with syllable changepoints (mean = 0.553 sec, mode = 0.233 sec) that best reflect the principal components changepoints (mean = 0.609 sec, mode = 0.200 sec). The fitting procedure yielded 52 syllables to explain 99% of the variance, which we truncated to 50.

For constructing the behavioral tensor, each vector of syllables over time is broken into bins of 9000 frames (5 min), and subsequently a third dimension of position bins using the *x*–*y* position of the mouse from extracted scalars. Position bins were constructed as ten concentric rings of equal area that capture the most frequently accessed portions of the video frame. In practical terms, the first 6 of these bins represent the floor of the arena, bins 7 and 8 capture a mouse’s rearing or exploratory activity against the wall, and the infrequently accessed bins 9 and 10 capture jumps and escape attempts. Crowd movies of each syllable were randomized and blinded and described by 2–4 experts for generating the labels of rear, pause, groom, and walk/run.

### Whole-brain Fos immunostaining with iDISCO+

After each mouse’s 45-min Fos-expression window following the 1-hr behavioral recording session, mice were anesthetized with Avertin and transcardially perfused with 20 ml of phosphate-buffered saline (PBS, Invitrogen AM9625) containing 1:1000 10 µg/ml heparin and subsequently 20 ml of ice-cold 4% paraformaldehyde (PFA). All harvested samples were post-fixed overnight at 4°C in 4% PFA in PBS, washed 3x in PBS the following morning and stored at 4°C for 4–7 days. Cohorts of 20 animals from one week of behavioral recordings were processed in parallel with the AdipoClear immunolabeling protocol, as detailed previously^64^. Samples were stained with 1:500 rabbit anti-Fos (Synaptic Systems, 226 003) and donkey anti-rabbit AlexaFluor 647 (ThermoFisher Scientific). All antibodies were centrifuged for 30 min at 12,000 rpm and 20°C before incubation. Primary antibodies for the complete set of 190 animals were batched together and then pre-measured and aliquoted at the start of the data collection to reduce technical variability in staining.

### iDISCO+ Imaging

At least 24 hours after clearing, iDISCO+ samples were imaged on a light-sheet microscope (Ultramicroscope II, LaVision Biotec) equipped with a sCMOS camera (Andor Neo) and a 2×/0.5 NA objective lens (MVPLAPO 2×) with a 6-mm working distance dipping cap. The imaging media was dibenzyl ether. Version v351 of the Imspector Microscope controller software was used. We imaged using 488-nm and 640-nm lasers. The samples were mounted horizontally, illuminated with a single sheet through the right hemisphere cortex, and scanned from dorsal to ventral with a step-size of 3 μm using the continuous light-sheet scanning method with the included contrast adaptive algorithm for the 640-nm channel (20 acquisitions per plane, max lightsheet NA), and without dynamic focusing for the 488-nm autofluorescence. In the *x*–*y* plane, voxels are 4.0625 μm × 4.0625 μm, yielding a field of view 10.4 mm anterior-posterior × 8.8 mm mediolateral. The small amount of sample shrinkage from the clearing protocol allowed the 6-mm working distance of the objective to capture the entire dorsal-ventral axis of the sample.

### Image processing and analysis

Registration to the Allen Reference Atlas CCFv3 and object segmentation were performed as previously described^64^. Briefly, the autofluorescent image acquired with the 488-nm laser was aligned to the reference atlas with a series of linear and non-linear transformations using the Elastix toolbox. These transformation parameters were then used to warp the coordinates of the Fos nuclei.

We used the TrailMap pipeline^64^ for segmenting the Fos nuclei from the stained images. While the architecture was unchanged from the previously published axon-detecting scenario, we trained a new model for detecting Fos signal by manually annotating 105 small, cropped volumes of nuclei selected to represent the variance in both signal and noise across the brain and across sample batches. The resultant probability maps were smoothed by convolving with a spherical nucleus-sized kernel and then processed for local maxima detection. Fos intensities for each detected maxima were calculated as the average of the raw image intensity values from a 6-connected neighborhood around the detected centroid.

Batch effects manifested as global shifts in image intensity, both of the detected Fos nuclei and of the background signal in the same channel. By detecting and proportionally scaling this global parameter before segmentation, two batches of 20 animals with reductions in intensity each were aligned with the average batch. However, one batch of 20 animals displayed intensity values too far from the average of the remaining samples and was excluded from further analyses, thus reducing the animals to n = 168.

### Tensor decomposition for behavioral data

Let *X* ∈ ℕ^*M*×*N*×*P*×*S*^ denote a four-dimensional tensor of non-negative counts *x*_*m,n,p,s*_ for each mouse *m* = 1, …, *M*, epoch *n* = 1, …, *N*, position bin *p* = 1, …, *P*, and behavioral syllable *s* = 1, …, *S*. We model the tensor as a weighted combination of factors,

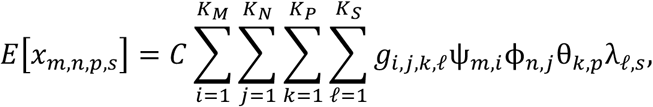

or, more compactly,

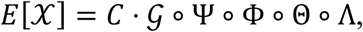

where *C* is a non-negative constant, *G ∈ R^K_M_×K_N_×K_P_×K_S_^* is the core tensor, Ψ ∈ *R^M×K_M_^* is a matrix with rows ψ_*m*_, Φ ∈ *R^N×K_N_^* is a matrix with rows ϕ_*n*_, Θ ∈ *R^P×K_P_^* is a matrix with columns θ_*k*_, Λ ∈ *R^S×K_S_^* is a matrix with columns λ_*l*_, and ∘ denotes a tensor-matrix multiplication. We recognize this as a 4-dimensional Tucker decomposition^42,43^. We impose non-negativity constraints on the factors so that the model can be interpreted as an additive combination of factors. Furthermore, we assume the factors are normalized so that they can be interpreted as distributions over positions, syllables, etc., which we encode via symmetric Dirichlet priors.

We fit the model using *maximum a posteriori* (MAP) estimation. We developed an expectation-maximization (EM) algorithm based on a Poisson data augmentation scheme. After augmentation, each factor admits a closed form coordinate update. The updates are naturally parallelizable. We wrote custom Python software using JAX for compilation and GPU acceleration. The software is available at https://github.com/lindermanlab/dirichlet-tucker. Mathematical derivations and further detail for the EM algorithm can be found in the package documentation.

### Hyperparameter selection for tensor decomposition model

The main hyperparameters to be determined are the number of factors, *K_M_, K_N_, K_P_*, and *K_S_*. We choose these parameters using cross-validation using a random, speckled test set. Specifically, we hold out a random subset of faces *X*_*m,n*_ from the data, i.e., we mask a random subset of (mouse, epoch) pairs. That way, we still have enough observed data to estimate the mouse loadings, ψ_*m*_, for each mouse, and the epoch loadings, ϕ_*n*_, for each epoch.

We evaluate the log likelihood of the held-out data using the parameters estimated from the training data. Rather than reporting the raw log likelihood, we compute the fraction of deviance explained, which is defined as,

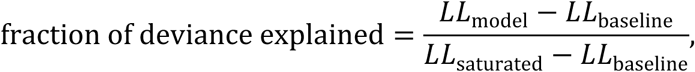

fraction of deviance explained where *LL*_model_ is the held-out log likelihood under the Tucker decomposition model, *LL*_*baseline*_ is the held-out log likelihood under a baseline of guessing the average behavior, and *LL*_saturated_is the held-out log likelihood under an “oracle” model that predicts that actual test behavior perfectly. The fraction of deviance explained is analogous to the fraction of variance explained in a linear Gaussian model.

We sweep over a four-dimensional grid of numbers of factors *K_M_* ∈ {2,4, …, 24}, *K_N_* ∈ {2,4, …, 8}, *K_P_* ∈ {2,4, …, 8}, and *K_S_* ∈ {2,4, …, 24}. The bounds of this search space were chosen manually to ensure that higher held out log likelihood could not be achieved with a larger model.

### Semi-nonnegative matrix factorization (semiNMF) of Fos data

The Fos data consists of cell counts ✔ ∈ ℕ^*M*×*v*^ and average intensities *y* ∈ ℝ^*M*×*v*^ for the *M* mice and *v* voxels. We model the cell counts for each mouse and voxel as Poisson random variables, *c*_*m,v*_ ∼ Pocλ_*m,v*_d, and we model the average intensity as conditionally Gaussian, *y_m,v_* ∼ N(μ_*m,v*_, σ^2^_v_/*c*_*m,v*_ d. Since the average intensity is computed from varying number of cells in each mouse and voxel, we rescale the conditional variance accordingly.

We model the mean count and intensity as a weighted combination of factors,

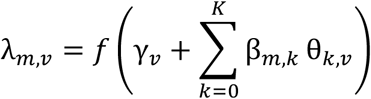

Where γ_*v*_ is a per-voxel effect, β_*m*_ ∈ *R^K^*^21^ are the per-mouse weights, and θ_*k*_ ∈ Δ^*v*^ are shared factors. The factors are non-negative and normalized to sum to one, so that they can be viewed as distributions over which voxels are active. The first factor is assumed to be constant, θ_0_ = 1_*v*_/*v* so that the first weight, β_0_, can be interpreted as a per-mouse intercept. The rectifying nonlinear function *f*(*x*) = *log*(1 + *e*^*x*^) is the “softplus” function, which ensures the expected cell count is non-negative.

The average intensities are modeled analogously,

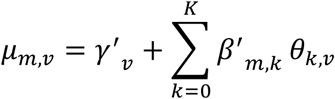

With their own per-voxel effects, γ^’^_*v*_, and per-mouse weights, β^’^_*m*_. The factors, θ_*k*_, are shared with the count model above.

We place a sparsity-inducing Laplace prior with rate α on the per-mouse weights for both counts and average intensity (excluding the per-mouse intercept terms). We place a uniform prior on the shared factors. We place a weakly informative, conjugate, inverse gamma prior on the per-voxel variance σ^2^_v_.

We fit the model using *maximum a posteriori* (MAP) estimation. We derived a coordinate ascent algorithm that cycles between updating the per-mouse weights, the shared factors, and the per-voxel variances and effects. The Poisson likelihood for counts renders the model non-conjugate. We follow standard practice for sparse generalized linear models (GLMs)^68^ and employ a proximal Newton update, which is based on a quadratic approximation of the log likelihood. To account for the sparsity inducing, Laplace prior, the proximal operator employs an inner loop of coordinate ascent to update each weight, β_*m,k*_, one at a time, holding the rest fixed. We use a similar proximal operator to enforce the non-negativity and normalization constraints for the shared factors.

The per-mouse weight updates can be parallelized across mice; likewise, the shared factor updates can be parallelized across voxels. We initialized the model using a heuristic, non-negative singular value decomposition (NNSVD) approach.

We wrote custom Python software using JAX for compilation and GPU acceleration, capitalizing on the parallelizability of the algorithm. The software is available at https://github.com/lindermanlab/fos. Mathematical derivations and further detail on the algorithm can be found in the package documentation.

### Hyperparameter selection for Fos semiNMF model

The main hyperparameters to be determined are the number of factors, *K*, and the sparsity parameter α. We choose these parameters using cross-validation using a random, speckled test set. Specifically, we hold out random, spatially contiguous blocks of voxels for each mouse. That way, we still have enough observed data to estimate the per-mouse weights, β_*m*_ and β^I^_*m*_, and the shared factors, θ_*k*_.

### Drug classification

We evaluated how well drug identity could be predicted from the per-mouse weights of the Fos and behavior models by training multiclass logistic regression models to predict the drug labels, *z*_*m*_ ∈ {0,1}^1G^ (a one-hot encoding of which drug mouse *m* was given), given the per-mouse weights (β_*m*_, β^’^_m_) of the Fos model or ψ for the behavior model. We used L2 regularization and selected the regularization strength using 5-fold cross validation over a grid of 31 values logarithmically spaced between 1e-15 and 1e15. We standardized each dimension of the regressors before fitting since we are using regularization. We fit the models using Scikit Learn (sklearn.linear_model.LogisticRegression and sklearn.model_selection.GridSearchCV).

We sorted the factors based on their contribution to drug classification, as measured by the magnitude of the drop in classification accuracy when each factor was left out. The first factor is the one that causes the largest drop in accuracy when omitted from the classifier.

### Fos-to-behavior prediction

After fitting the Tucker decomposition of behavior data and the semiNMF model for Fos data, we trained linear models to predict the per-mouse weights of one model given the weights of the other. Since the per-mouse weights of the behavior model are non-negative and normalized, we clipped and normalized the output of the linear model as follows,

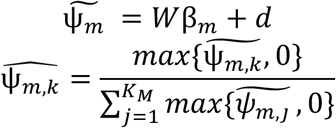

Where *W* ∈ *R^K_M_×K^* are the weights of the linear model and *d* ∈ *R^K_M_^* is the intercept. We trained the linear model on 4 folds of data, each with a random subset of 75% of the mice, and we evaluated predictions on the held-out 25% for each fold.

We also constructed a “hybrid model” to test if predictions could be improved by incorporating drug identity by incorporating an indicator into the model,

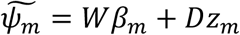

where *D* ∈ *R^K_M_×1G^* is a matrix of intercepts for each drug, and *z*_*m*_ ∈ {0,1}^1G^ is a one-hot indicator of which drug mouse *m* was given. (The intercept *d* is redundant in this model formulation.) We train the model using the true drug labels, and on test data we replace *z*_*m*_ with *z*Ü_*m*_, the predicted probabilities output by the drug classifier trained on Fos weights, β_*m*_.

For end-to-end predictions, we started with the saline average Fos measurements and constructed a synthetic Fos map by modulating the Fos cell count, *c*_*m,v*_, across. We considered two types of spatial perturbations: up-regulating all voxels in a region, and “correlated whole-brain modulation” in which we perturbed voxels according to their correlation with a seed voxel. Let ρ_*uv*_ ∈ [−1,1] denote the correlation between seed voxel *u* and target voxel *v*, and let σ_*v*_ denote the standard deviation of voxel *v*. In correlated whole-brain modulation, we perturb each voxel *v* by an amount ρ_*uv*_σ_*v*_.

We used the semiNMF model with the learned factors, θ_*k*_, to estimate the most likely weights, β^ã^_*m*_, for the perturbed Fos maps. Then we map the estimated weights (and the predicted drugs, *z*Ü_*m*_, for the hybrid model), through the linear model to estimate weights of the behavior model. Finally, we combine the estimated behavior weights with the shared factors of the behavior decomposition model to construct a predicted behavior tensor.

### Behavior-to-Fos prediction

We perform the same analysis in reverse when predicting per-mouse weights β_*m*_ of the Fos model given per-mouse weights ψ_*m*_ of the behavior model. The only difference is that in this direction, we need to take extra care to account for batch effects in the Fos weights. The Fos data was collected in batches of 20 mice, and each batch differed slightly in the number of cells and average intensity due to differences in the immunostaining, etc. We account for these batch effects by including an extra term in the linear model, *Bb*_*m*_, where *B* is a matrix of per-batch intercepts and b_L_ is a one-hot indicator of which batch mouse *m* belongs to.

For the behavior-to-Fos prediction, we first constructed a synthetic tensor of syllable counts over space and time by starting with the saline average behavior and then artificially increasing the amount of pausing, grooming, rearing, or locomotion. Note that the total number of video frames per epoch is fixed, so an increase in one category of behavioral syllables necessarily decreases the others. We use the tensor decomposition model to estimate the weights, ψ^ã^_*m*_, corresponding to the perturbed behavior. Then, using the linear model, we predict the corresponding change in Fos weights. Finally, we map the change in Fos weights from the saline average to a predicted change in Fos measurements.

### Bootstrapping analysis

To assess the significance of our Fos to/from behavior analyses we used a bootstrap analysis to estimate confidence intervals of our predictions. We generated 250 replicate datasets using a stratified bootstrap approach: each drug group was resampled with replacement to produce replicate datasets with the same numbers of mice per drug group as in the original dataset. We trained the behavioral tensor decomposition model and Fos semiNMF model on the 250 bootstrap replicates, and we trained linear models to predict Fos weights from behavior weights and vice versa. Then, we computed the distribution of change in behavior for a given change in Fos activity across bootstrap replicates. The quantiles of this distribution provide an estimate of the confidence interval for the prediction.

## Data availability

Behavior data and whole-brain Fos data are freely available at https://dandiarchive.org/dandiset/001204

## Acknowledgements

We thank members of the Luo and Linderman labs for discussion, and Lijun Qi and Yunming Wu for comments on the manuscripts. This work was supported by NIH BRAIN Initiative grants R01-NS131987 (to L.L. and S.W.L.), R01-NS105698 (to L.L.), and fellowships from the McKnight and Sloan Foundations (to S.W.L.). L.L. is an HHMI Investigator.

## Author contributions

D.F., A.M, and L.L. designed the experiments with feedback from S.W.L. D.F. and A.M. collected all data. D.F., A.D., and N.T. preprocessed all imaging and behavior data and built the first versions of the models. X.G. and S.W.L. built the models in this paper. D.F., T.W., A.D., and S.W.L. contributed key analyses of the model outputs. J.H.S. and S.R.D. provided important insights for structuring our analyses. D.F., S.W.L., and L.L. wrote the paper.

## EXTENDED DATA

**Extended Data Fig. 1.**
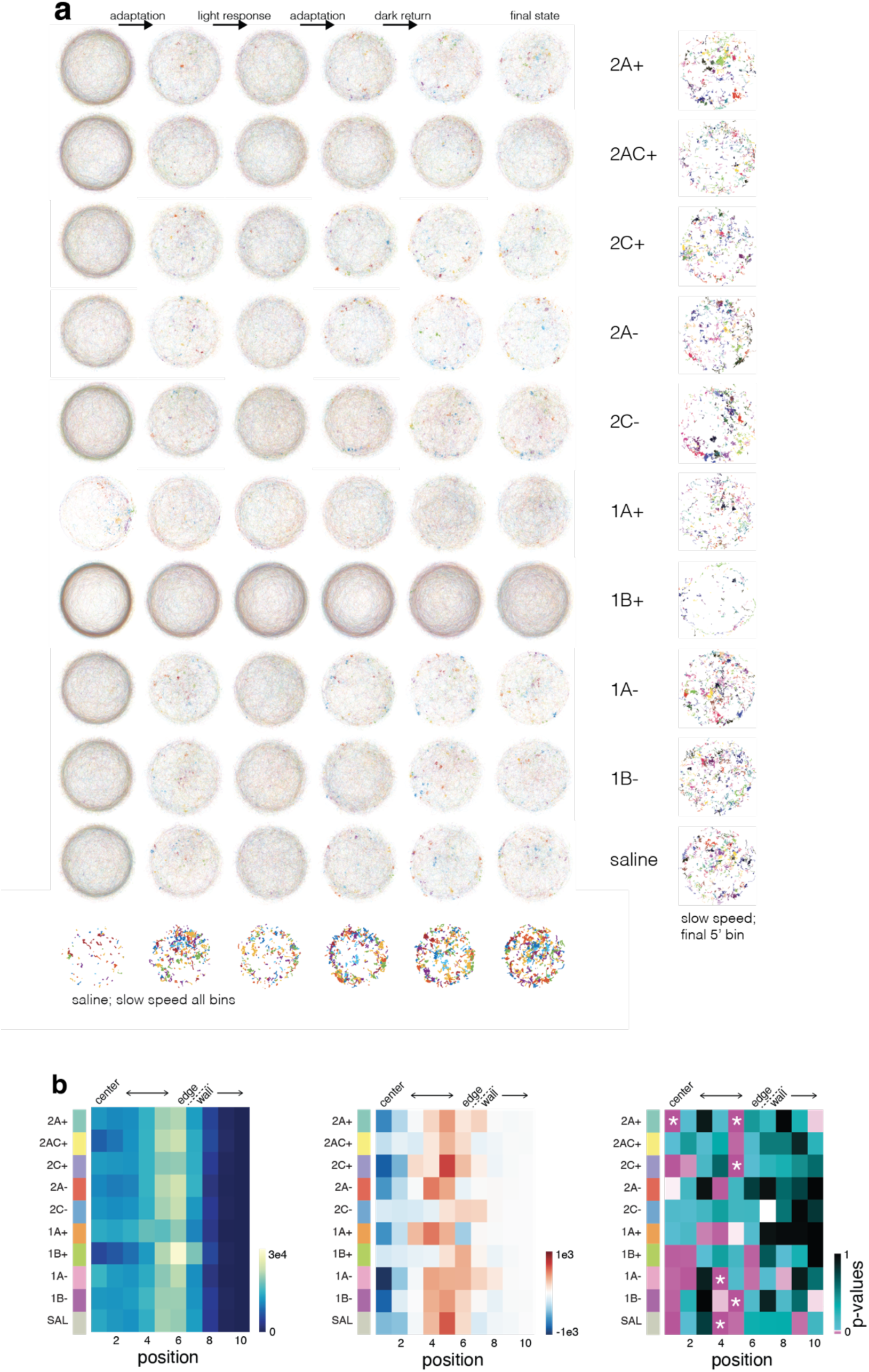
Additional data on the effect of drugs on spontaneous behavior measured by scalar. **a**, Tracklets for all drug treatment in 6 separate 5-min periods (0–5’, 26–30’, 31–35’, 41–45’, 46–50’, and 55’–60’) from left to right. Right-most column represents slow-speed tracklets in the final 5-min window as described in Fig. 1b. Bottom-most row represents slow-speed tracklets for each of the 6 visualized time bins for saline mice. **b**, Left, drug group average occupancy of 10 spatial bins defined as concentric rings of equal area. The edge of the floor is approximately between bins 6 and 7. Mice rearing and exploring the wall surfaces appear in bins 7–8 with jumping above bin 8. Rows sum to 1.08×10^5^, the number of frames in 60’. Center, differential occupancy of position bins in the 5 min before and after the LED onset. Rows sum to zero. Right, t-test for deviation from zero for the LED onset response; asterisks represent p-values passing a Bonferroni-corrected α for *m* = 10. 2AC+, 2C–, and 1A+ treated groups did not show a significant reduction in center occupancy and the 2C– group also did not show an increase in occupancy of the edge of the arena.

**Extended Data Fig. 2.**
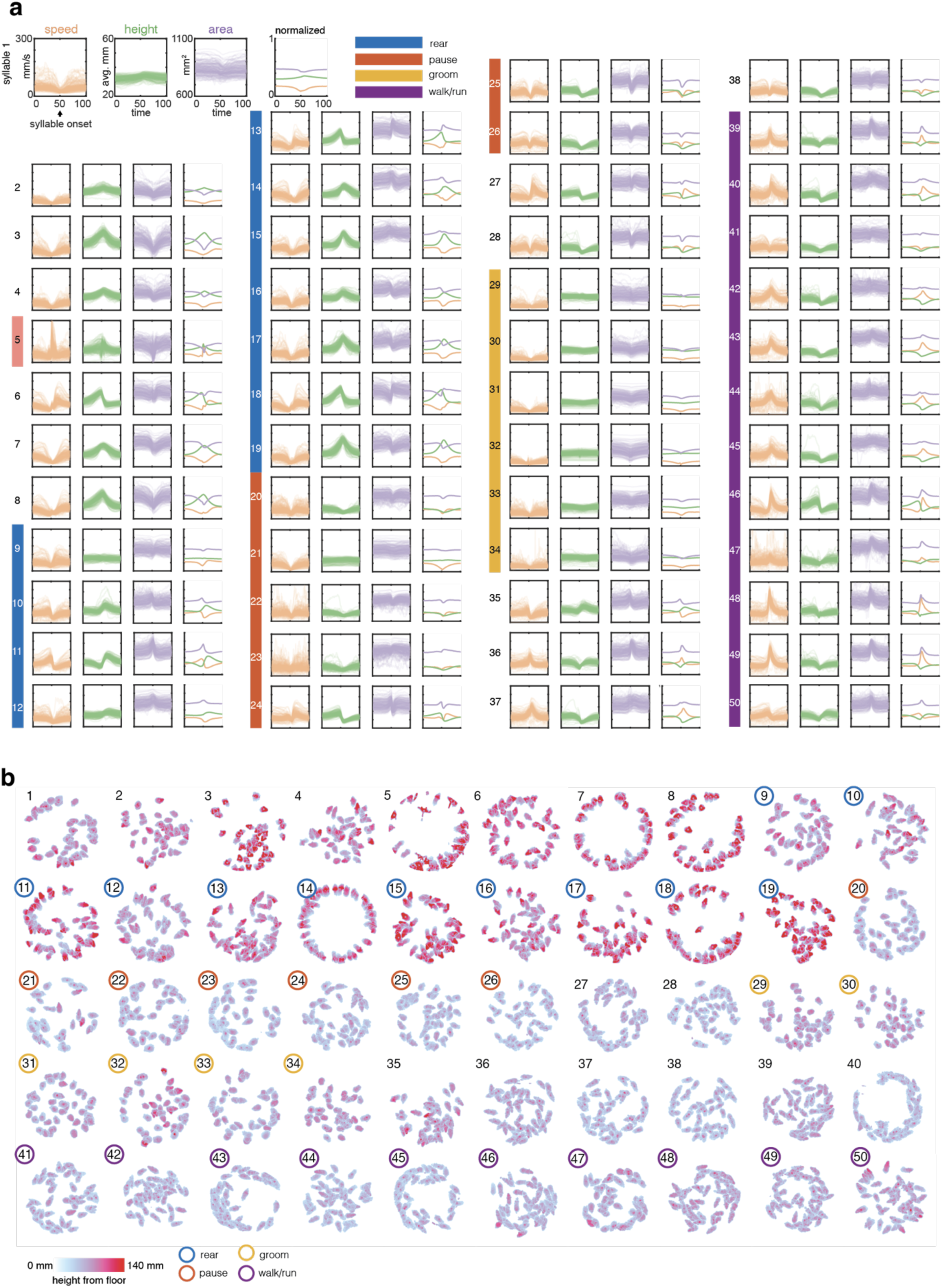
Additional data on the effect of drugs on spontaneous behavior as measured by MoSeq. **a**, Speed, height, and area plots for all 50 syllables. Time = 50 represents time-locked onset of the syllable. The four major behavioral clusters are highlights with colored bars. Syllable 5, the jumping syllable, is also highlighted. **b**, Single frame extracted from crowd movies of each individual syllable during the display of the syllable. Height of each individual pixel is color coded from the depth camera output and highlights the mouse body shape and position for syllables with high average height plots in **a**.

**Extended Data Fig. 3.**
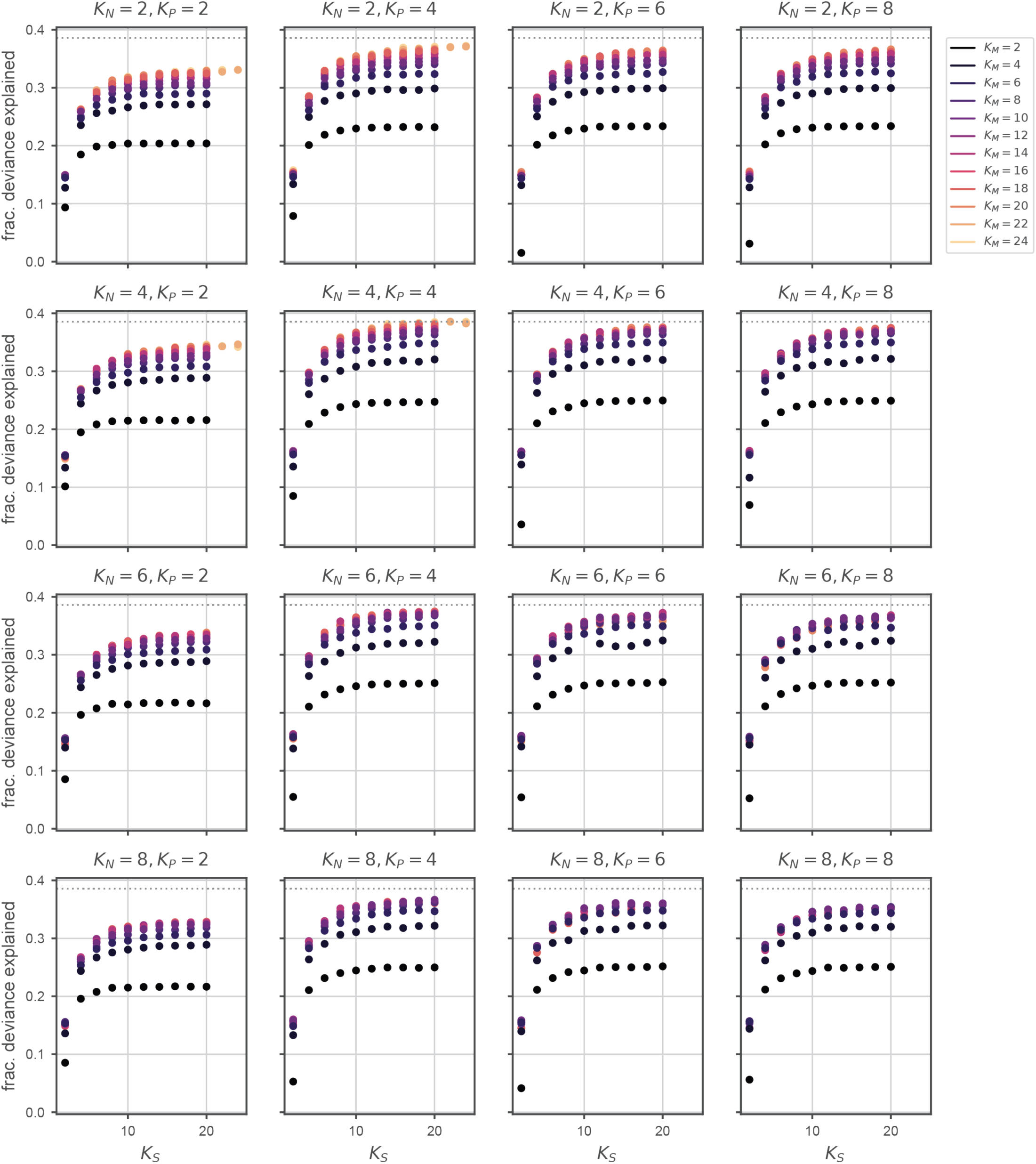
Hyperparameter selection for behavior model. The hyperparameters include the number of shared factors (and per-mouse weights) *K_M_*, the number of epoch factors *K_N_*, the number of position factors *K_P_*, and the number of syllable factors *K_S_*. The optimal setting of hyperparameters was chosen by a grid search, which identified the combination that yielded the highest fraction of deviance explained on held-out test data (Methods).

**Extended Data Fig. 4.**
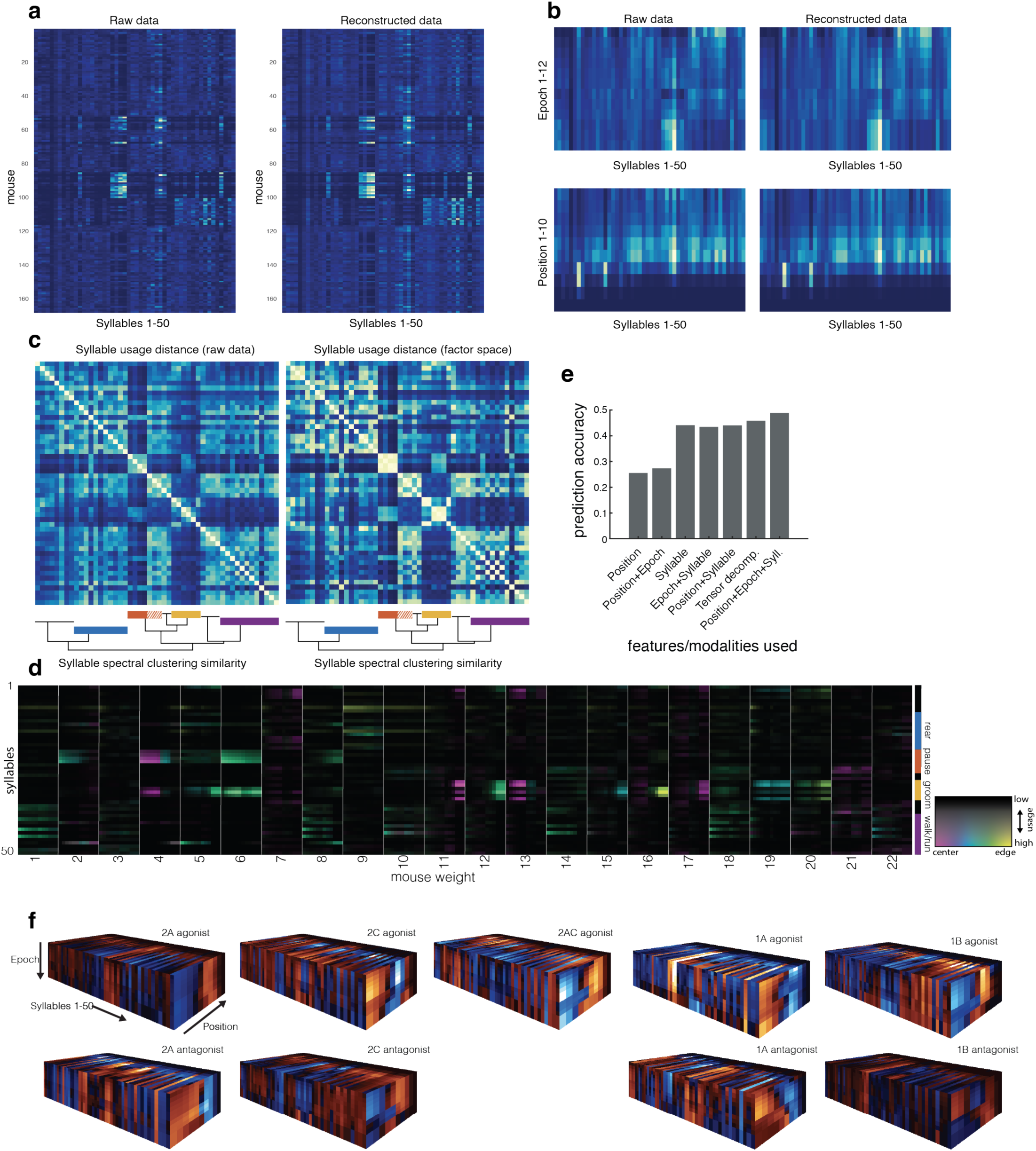
Decomposition of behavior. **a**, Side-by-side view of syllable usage by mouse from raw data (left, as in Fig. 1g) and reconstructed data from the Tucker decomposition (right). **b**, Side-by-side view of raw and reconstructed behavioral data from the behavioral tensor. Top, epoch by syllable; bottom, position by syllable. **c**, Cosine distance between individual syllable usage vectors across n = 168 mice is similarly captured by the reconstructed data. **d**, All 22 mouse factors from the decomposition shown as usage heatmaps of 50 individual syllables over 12 epochs. The intensity of the color corresponds to usage while the hue corresponds to the modal position bin (1–12) for that syllable/epoch pairing. **e**, Prediction accuracy for the drug classification multi-class logistic regression, using one or more modalities. For example, “Position” model predicts drug solely based on a mouse’s average position in the arena. “Tensor decomp” indicates performance of the logistic regression using the 22-dimensional mouse weights from the Tucker decomposition model, which is much lower-dimensional than the raw, 6000-dimensional position+epoch+syllable count vector. **f**, The weighted combination of shared factors using the weights of the logistic regression model show which epochs, syllables, and positions are most predictive of each drug. Expected increases in position+epoch+syllable representation are colored in orange and expected decreases are colored in blue. Elements of the tensor are summed and represented on each face.

**Extended Data Fig. 5.**
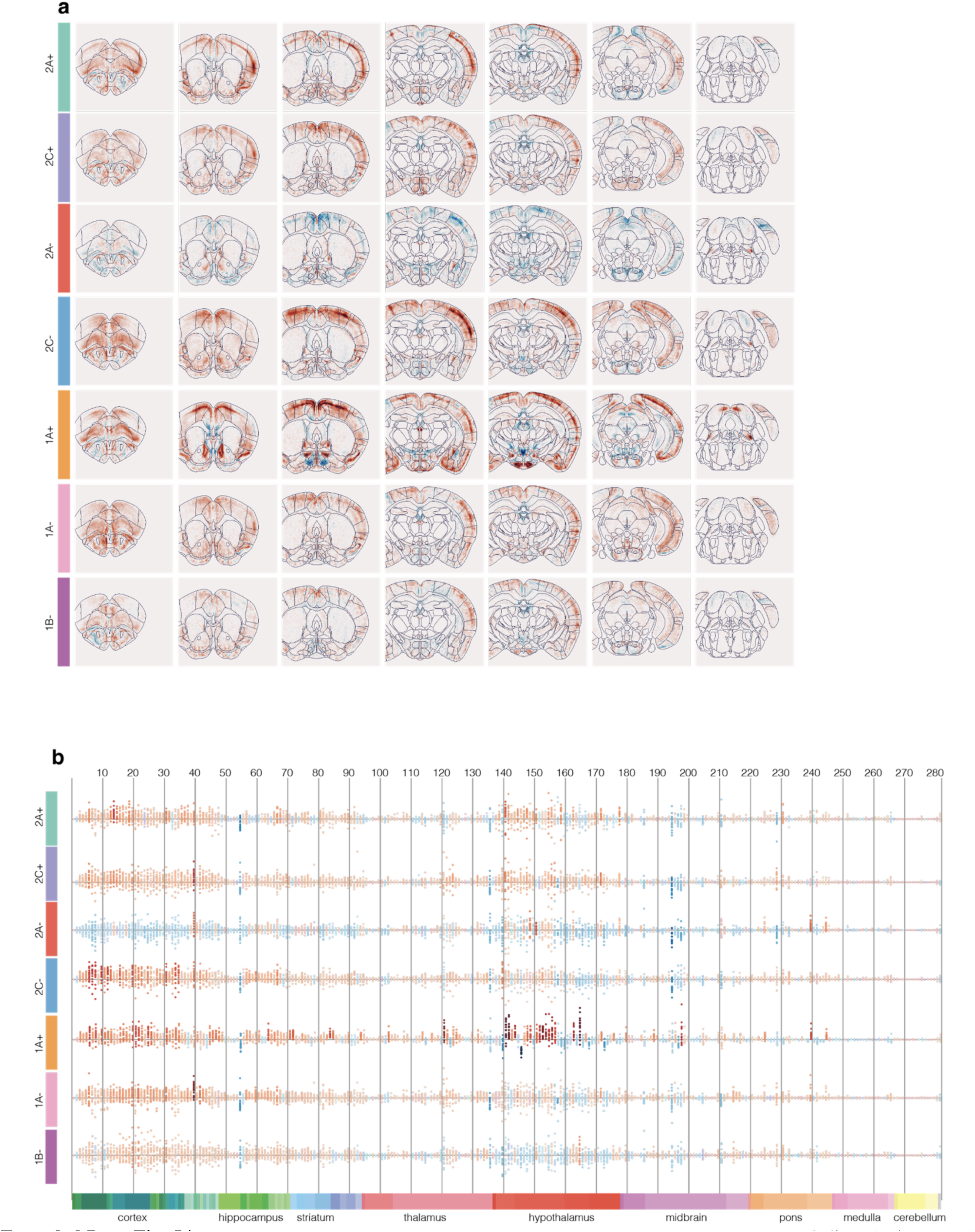
Additional data on drugs’ effect on whole-brain Fos map. a, 100-µm coronal slices (as in Fig. 3g) across anteroposterior axis for the 7 remaining drug treatment groups. b, Average saline-subtracted Fos intensity (as in Fig. 3h) for the 7 remaining drug treatment groups in a for all nuclei in each of 282 Allen-brain regions at the bottom, color coded according to Allen Brain Atlas^45^. Each dot represents data from an individual mouse. See Table S3 for the names of the enumerated regions.

**Extended Data Fig. 6.**
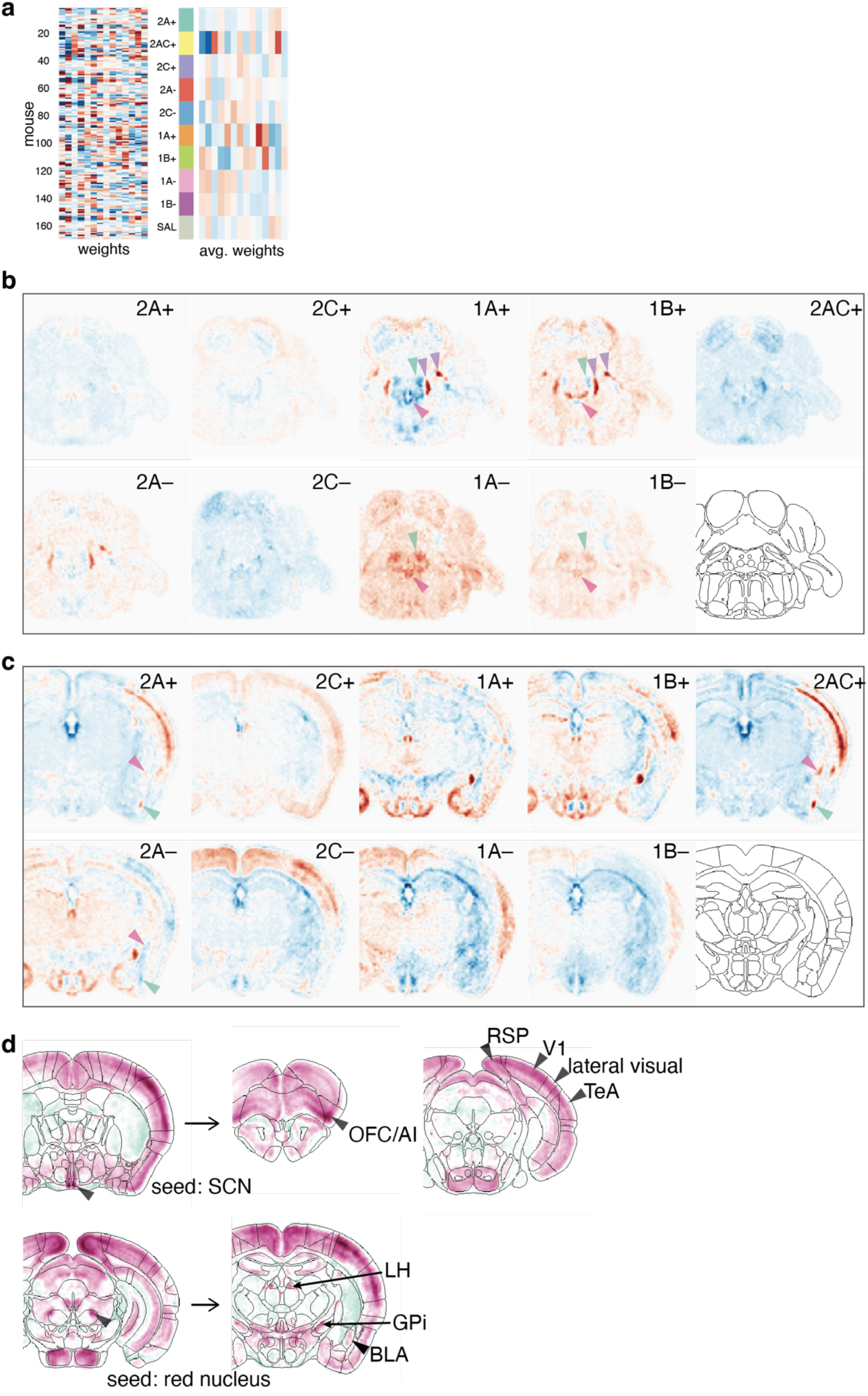
Additional data on decomposition of Fos map. **a**, The Fos intensity weights for each mouse (left) and the average for each drug group (right). **b**, A coronal section at the brainstem from prediction filtered factors (as in **Fig**. **5g**). Arrowheads mark PBN and LC (purple), Barrington’s nucleus (green), and nucleus incertus (pink). **c**, Data as in **b**, a coronal section highlighting the expected changes in amygdalar hotspots of 2A+ interacting drugs with a sign inversion to the expected change in 2A–. BLAv, green arrowhead, lateral amygdala, pink arrowhead. **d**, Voxel-wise correlation maps as in **Fig. 5j** for seed voxels in SCN (top) and red nucleus (bottom). Arrowheads mark notable hotspots of high positive correlation.

**Extended Data Fig. 7.**
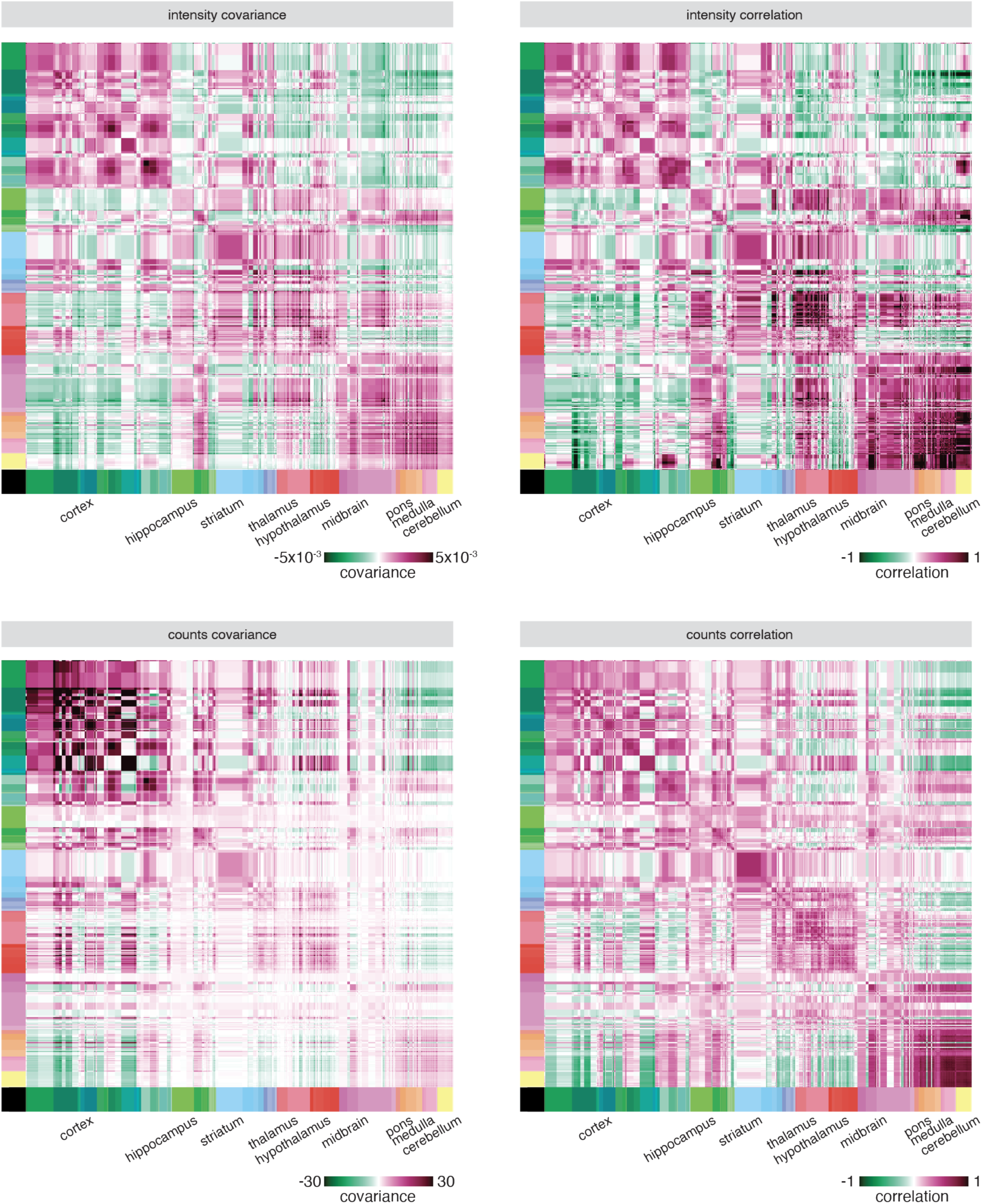
Voxel-wise whole-brain covariance and correlation maps based on Fos intensity and Fos+ cell counts. **a**, Brain-wide covariance map (as estimated by the 14-dimensional factor model) summarized at the level of Allen Brain Atlas regions. Regions are scaled proportionally to their volume and entries on the diagonal represent the mean covariance of the voxels within that region. Covariances are calculated on the voxel-wise transformed Fos intensities (**Methods**). Same as **Fig. 5i** for ease of comparisons with other panels. **b**, Brain-wide correlation map. Correlations calculated separately for each individual voxel and averaged within Allen Brain Atlas regions to populate the entries on the diagonal. Off-diagonal entries represent average correlation between each pair of voxels in each region. Calculated from Fos intensities as in (**a**). **c**, Brain-wide covariance map as in (**a**), calculated with voxel-wise counts of Fos+ cells (**Methods**). **d**, Brain-wide correlation map calculated as in (**b**), originating with voxel-wise Fos counts as in (**c**).

**Extended Data Fig. 8.**
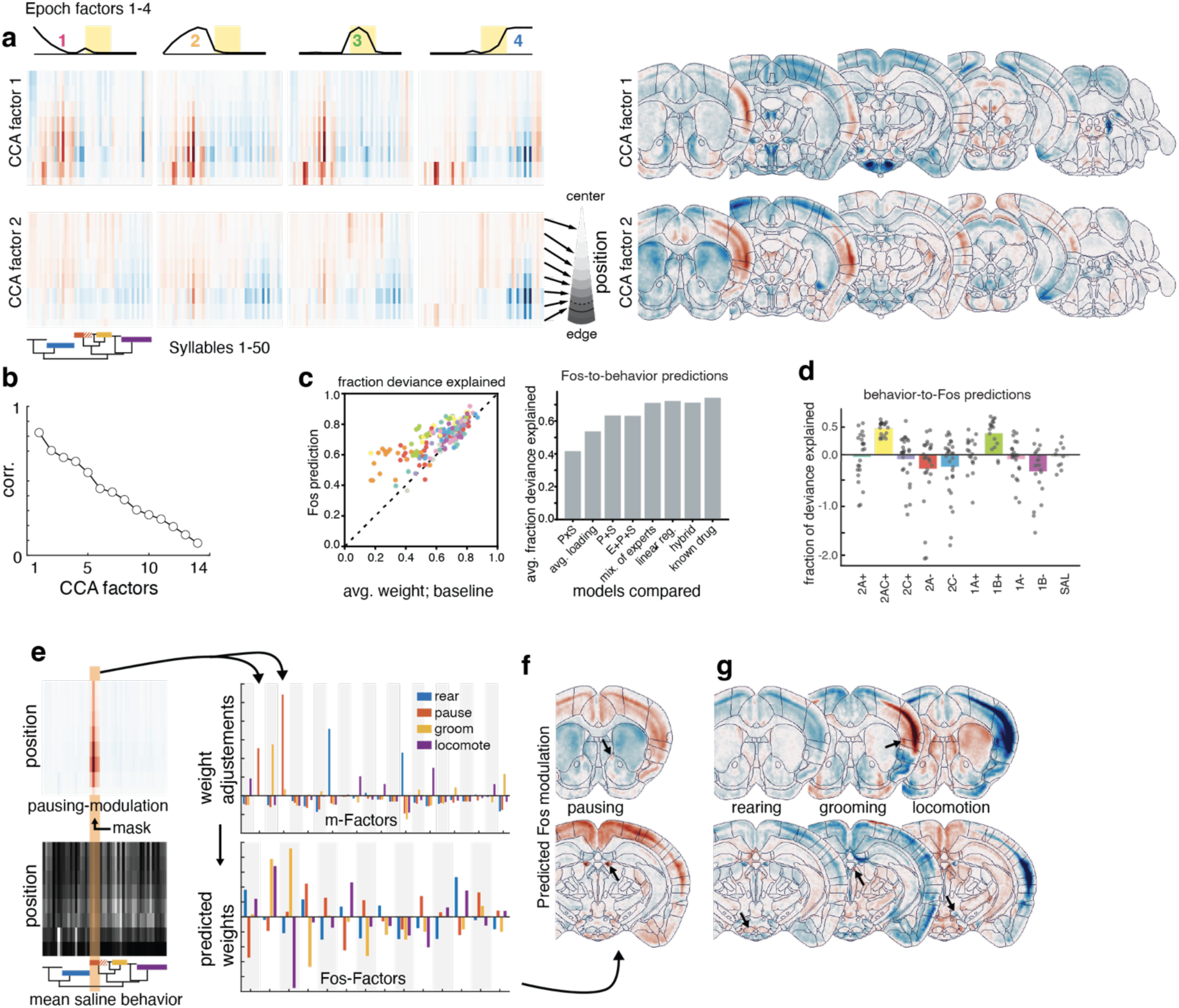
Linking behavior and Fos map using canonical correlation analysis (CCA) and linear regression. **a**, CCA generated 14 canonical factors, the first three of which are shown here. Factors are in both behavior space (left, represented as syllable × position heatmaps across the 4 epochs; values are expected difference from saline) and brain space (right, as expected difference from saline). **b**, Correlation values for all 14 canonical factors, the first of which is 82% correlated between the two modalities. The first 5 factors are over 50% correlated. **c**, Left, individual mouse data summarized in **Fig. 6b** for Fos-to-behavior prediction (see **Methods**). Right, fraction of deviance explained in the Fos-to-behavior prediction across models selected with varying quantities of information. P×S: using the marginal distributions of position and syllable, separately; avg. loading: using the average weight in the tensor decomposition model; P+S: using the marginal distribution of position and syllable, jointly; E+P+S: using the marginal distribution of position and syllable, jointly for each epoch; mix of experts: combination of drug group average behavior weighted by predicted drug group probability given Fos; linear reg: predicts behavior weights given Fos weights; hybrid: combines mix of experts and linear regression but overfits slightly; known drug: using average behavior of true drug group, which is not known to other models. Linear regression, our selected model, is second only to the inclusion of drug identity. **d**, Predictive power of behavior-to-Fos linear models depicted as fraction of deviance explained for held out test mice, relative to a baseline model of predicting average behavior. Predicting Fos from behavior could be more challenging because of batch effects in Fos intensity, which manifest in the Fos weights even though the factor model attempts to capture them with a batch covariate. Also, neural activity measured in Fos may simply contain information that is not present in behavioral syllables. **e**, Predicting Fos maps from synthetic modulations on average saline behaviors. Left, upmodulating pausing syllable usage from the saline average behaviors (gray) yields an artificial tensor that has extra pausing (red). The behavioral factor weights for this artificial pausing tensor are predicted (orange bars only, increased weights on factors 2 and 4), then using the coefficients from the linear regression, the Fos factor weights are predicted and combined with all 14 Fos factors to generate predicted brain maps. **f**, Values represent expected change in Fos from a similarly predicted saline average map without modulation; two coronal planes are shown. **g**, Similarly produced maps for rearing, grooming, and locomotion, following the same synthetic strategy of upmodulating groups of syllables across all positions and epochs.

**Extended Data Fig. 9.**
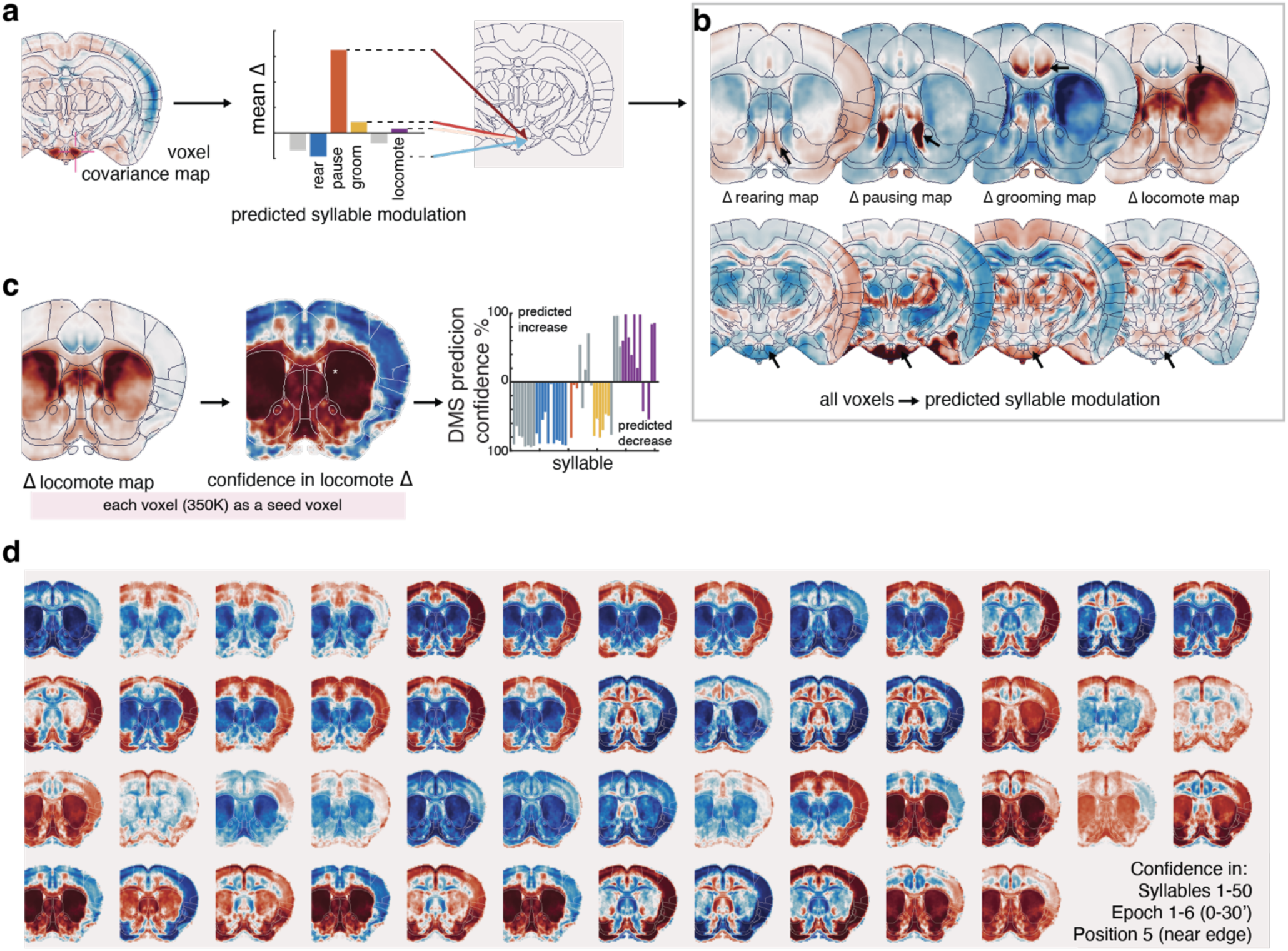
Linking behavior and Fos map via correlated whole-brain modulation. **a**, Voxel-wise correlation map for a seed voxel in ventral premammillary nucleus (PMv), as in **Fig. 6d**, is used as a perturbation map for predicting changes in behavior. Starting with the average saline brain, 10 artificial Fos+ cells are added to the seed voxel (average of 23 cells per voxel across all voxels in all brains) and up to 10 artificial cells are added to or subtracted from all other voxels, weighted by each voxel’s correlation (–1 to 1) with the seed. Predicting behavior (following a procedure reverse to what is described in **Extended Data Fig. 8e** in the behavior-to-Fos direction) generated the bar plot to the right. In this example, voxels in the hotspot in PMv (at left) will have high scores for pausing behavior, low positive scores for grooming, and negative scores for rearing. **b**, Shown here are two coronal planes of maps representing the four main behavioral categories in brain-space. Note that these maps are not imputed Fos values at each voxel; rather, these values represent the predicted change in each behavior that follows from a standardized perturbation originating at each voxel. By generating a map wherein these seed voxels contain the values of the predicted changes in behavior, we can visualize the extent of the hotspot in PMv that predicts pausing behavior and other behaviors as well. Repeating this procedure at every seed voxel in the brain yields complete maps of brain regions that predict high magnitude behavioral modulation for each specific subset of behavior. **c**, Left, the map of all seed voxels whose perturbation predicts modulation of locomotion behavior. Large positive values in dorsomedial striatum indicate that synthetically high Fos counts in this region (and the brain-wide Fos pattern that covaries with it) predict increased locomotion in the arena. Center, after testing each seed voxel’s prediction across bootstrapped model fits, we generated a voxel-wise map of the percentage of the fits at each voxel that shows a positive (red) or negative (blue) modulation of locomotion. Values approaching 1.0 indicate high confidence that perturbations at that voxel predict increases in locomotion, but this does not carry information about the magnitude of the predicted change. Right, as confidence scores from each seed voxel can be generated for each syllable+epoch+position triplet, we averaged the confidence scores for predictions of all 50 syllables (in epochs 1–6, position bin 5) following perturbations at each seed in a 27-voxel cube at the asterisk in the previous map. Perturbations in this region confidently predict increases in most locomotion syllables and confidently predict decreases in grooming and rearing syllables. **d**, Confidence maps, as in **c**, for each of the 50 syllables, averaged across epoch 1–6, position bin 5. Values in a given voxel approaching 1.0 and –1.0 indicate high confidence that a perturbation originating at that voxel will well-predict modulation of that specific syllable at the edge of the arena and in the first 30 min.

---

**Table S1:** Description of 50 MoSeq syllables.

| Order in Hierarchical Sort Ranking | Group at h=1 | Annotation |
| --- | --- | --- |
| 1 | 2 | compact body, initiate nose lift, XY still, height flat, area unchanging |
| 2 | 2 | scrunch or groom or small rear, XY still or slow, area small |
| 3 | 2 | unsupported rear / standing, area small |
| 4 | 2 | mixed syllable, a little rear or back arch during a groom, area small |
| 5 | 2 | jump |
| 6 | 2 | fall from rear to a compact position, then extend and walk, XY double peak |
| 7 | 2 | mid-height rear towards wall edge |
| 8 | 2 | rear towards wall, followed by a turn of the head |
| 9 | 4 | lift head up, turn of the head, XY paused + slow before/after, height unchanging |
| 10 | 4 | pause and look up, slight decrease in area |
| 11 | 7 | decelerate and initiate rear, XY still |
| 12 | 7 | nose up / low rear, XY slow |
| 13 | 7 | middle of rear / fall from rear |
| 14 | 7 | supported rear (facing wall), big area |
| 15 | 7 | maintain / steady rear, big area |
| 16 | 7 | head lift and look |
| 17 | 7 | mid-height rear |
| 18 | 7 | fall from rear, right turn |
| 19 | 7 | peak of rear |
| 20 | 5 | pause if in motion, compact shape / little scrunch, XY still, area unchanging |
| 21 | 5 | completely still, height flat, area unchanging |
| 22 | 5 | a slow walking scrunch, nose tuck, XY slow with an additional dip in speed |
| 23 | 5 | slow walking or still, looks down, nose forward, XY slow, area unchanging |
| 24 | 5 | fall from rear to a compact position, then extend and walk, XY motion from fall only |
| 25 | 5 | scrunch |
| 26 | 5 | pause between walks |
| 27 | 1 | turn left and walk |
| 28 | 1 | pause and scrunch and go, height unchanging |
| 29 | 1 | groom, itch, using front legs, XY stationary before/during/after, height unchanging, area small/unchanging |
| 30 | 1 | groom, height flat, area small/unchanging |
| 31 | 1 | groom, area small/unchanging |
| 32 | 1 | pause/scrunch without nose fully tucked, height flat, area small |
| 33 | 1 | nose tuck, XY still, height flat, area unchanging |
| 34 | 1 | groom, height flat, area small/unchanging |
| 35 | 6 | low rear, head exploration, XY varied but slow |
| 36 | 6 | dart / hop forward / lurch, then decelerate, XY fast, area briefly increases |
| 37 | 6 | a full run sequence from acceleration to stop, XY fast |
| 38 | 6 | decelerate / look down |
| 39 | 3 | extend nose out in front + forward run, big area increases |
| 40 | 3 | counterclockwise wall run, big area |
| 41 | 3 | initiate a slow step forward, XY slow + brief speed dip before start |
| 42 | 3 | initiate and sustain a run, XY fast, area unchanging |
| 43 | 3 | clockwise wall run, big area |
| 44 | 3 | the second half of a short run, XY finishes peaking then decelerates |
| 45 | 3 | an acceleration from any starting speed, big area |
| 46 | 3 | fall from rear, initiate dart, then decelerate fast, big area increases |
| 47 | 3 | walk forward, big area |
| 48 | 3 | initiate and sustain a fast run, area sharply increases |
| 49 | 3 | dart / lurch, area increases |
| 50 | 3 | forward walk, area increases |

**Table S2:** Possible interpretations of agonist and antagonist effects.

| Fos intensity | Example (in **Fig. 3f**) | Possible interpretations<sup>1</sup> |
| --- | --- | --- |
| Enhanced by agonist; suppressed by antagonist<sup>2</sup> | 2A in somatosensory cortex | Receptor is expressed and activated in saline control by the open field behavior at an intermediate level |
| Modulated by agonist; no effect by antagonist | 1B in BNST or dorsal striatum (Fig. 3e) | Receptor is expressed but not substantially activated in control by the behavior |
| No effect by agonist, modulated by antagonist | 2A or 2C in BNST | Receptor is activated at a saturation level by the behavior in control, so only downregulation has an effect |
| Agonist and antagonist have the same effect | 2C in somatosensory cortex | Receptor is expressed in mutually connected excitatory and inhibitory neurons in the same region, and agonist and antagonist preferentially act on receptors in either of the populations |
| Neither agonist nor antagonist has an effect | 2A or 2C in dorsal striatum | Receptor is not substantially expressed in the brain region of interest |
| <sup>1</sup> We list only one of several possible interpretations. One complicating factor is that serotonin can act on receptors expressed on presynaptic terminals of neurons far away from the region of interest.<br><sup>2</sup> The opposing effects can be the other way around (enhanced by antagonist, suppressed by agonist). The simplest interpretation is that activation of the receptor suppresses neuronal activity (as is the case for 1A and 1B receptors, which are coupled to G<sub>i</sub> and known to suppress neuronal activity). The other interpretation is that the receptor activates local inhibitory neurons, which in turn suppress the activity of nearby neurons. |  |  |

**Table S3:**
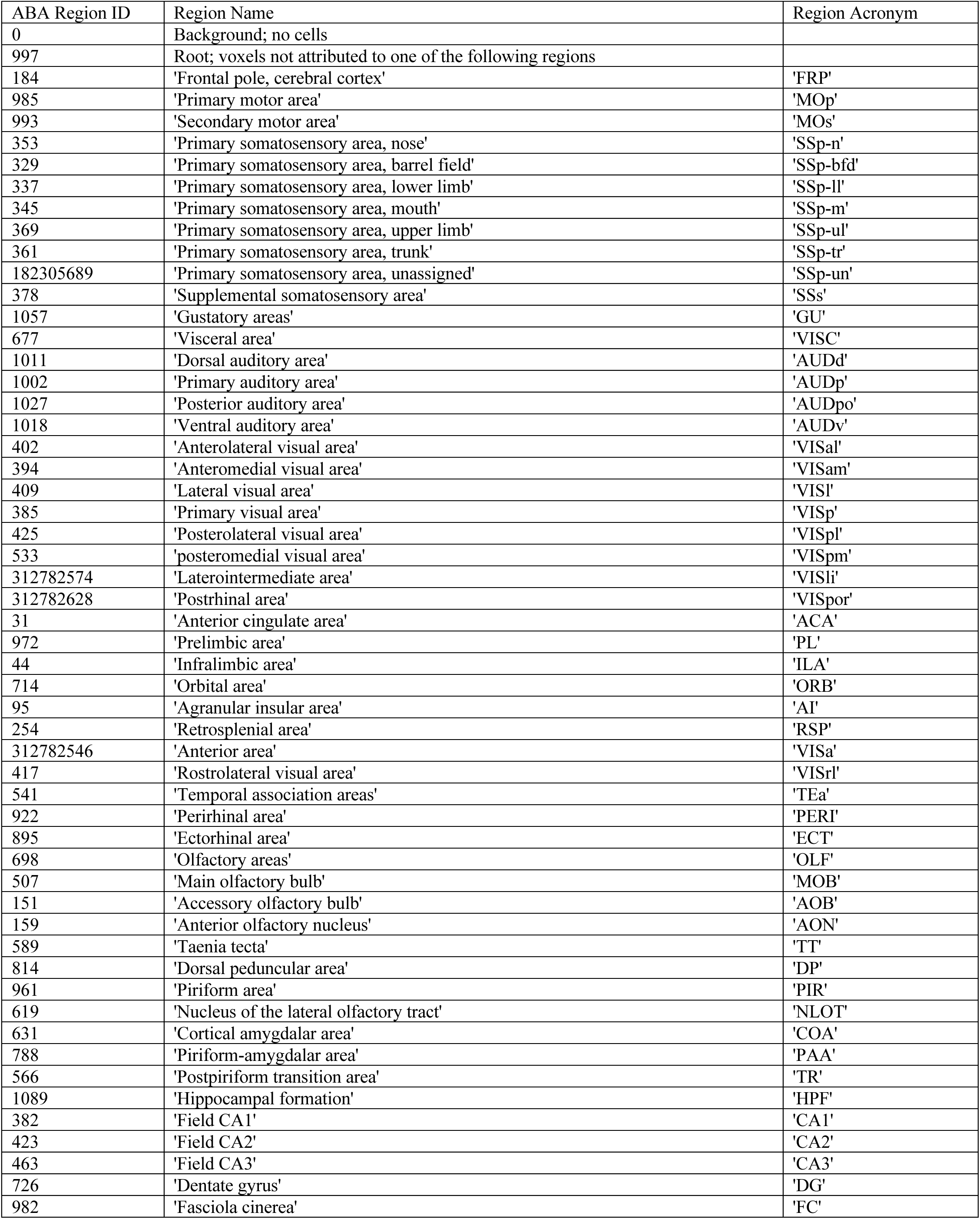
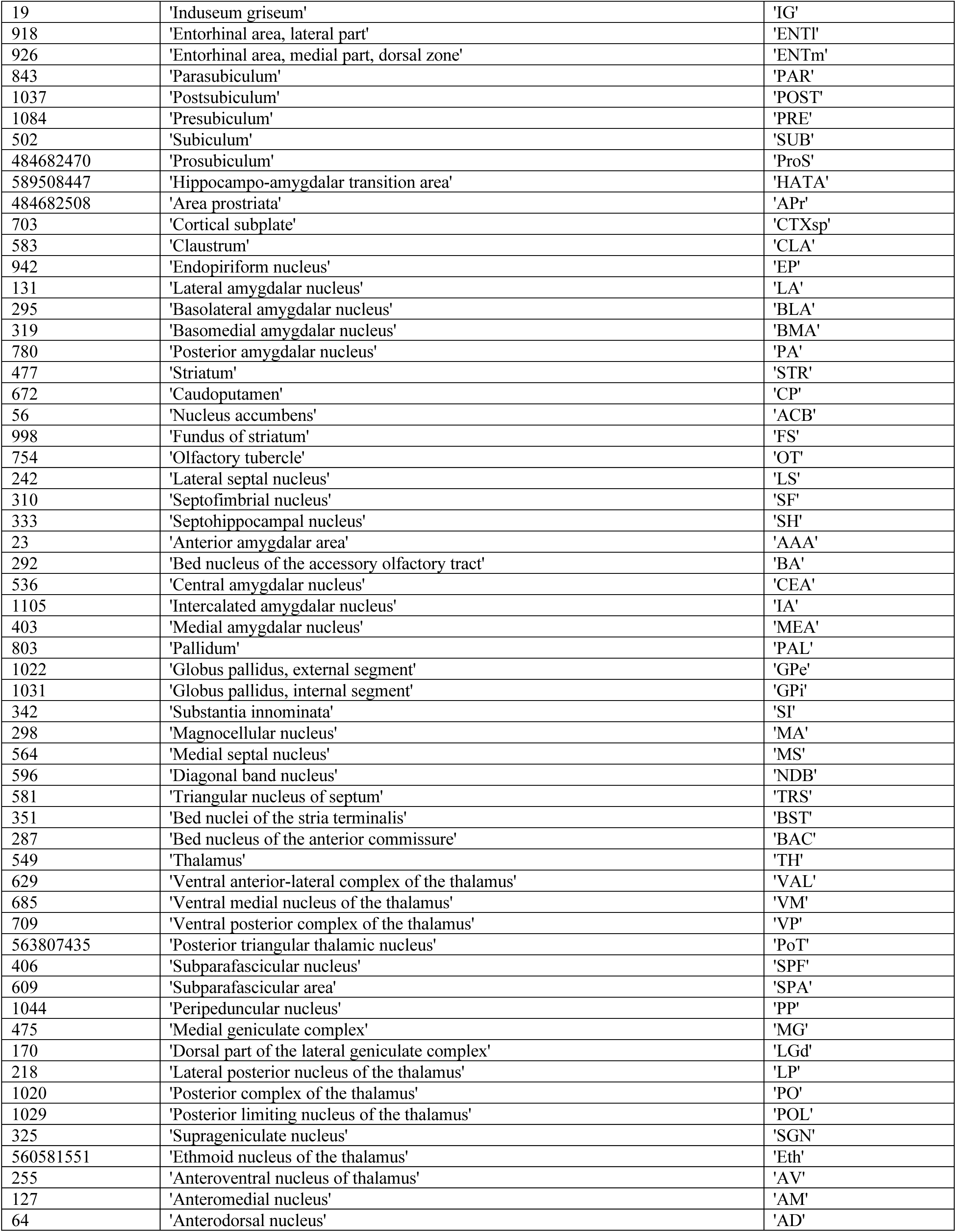
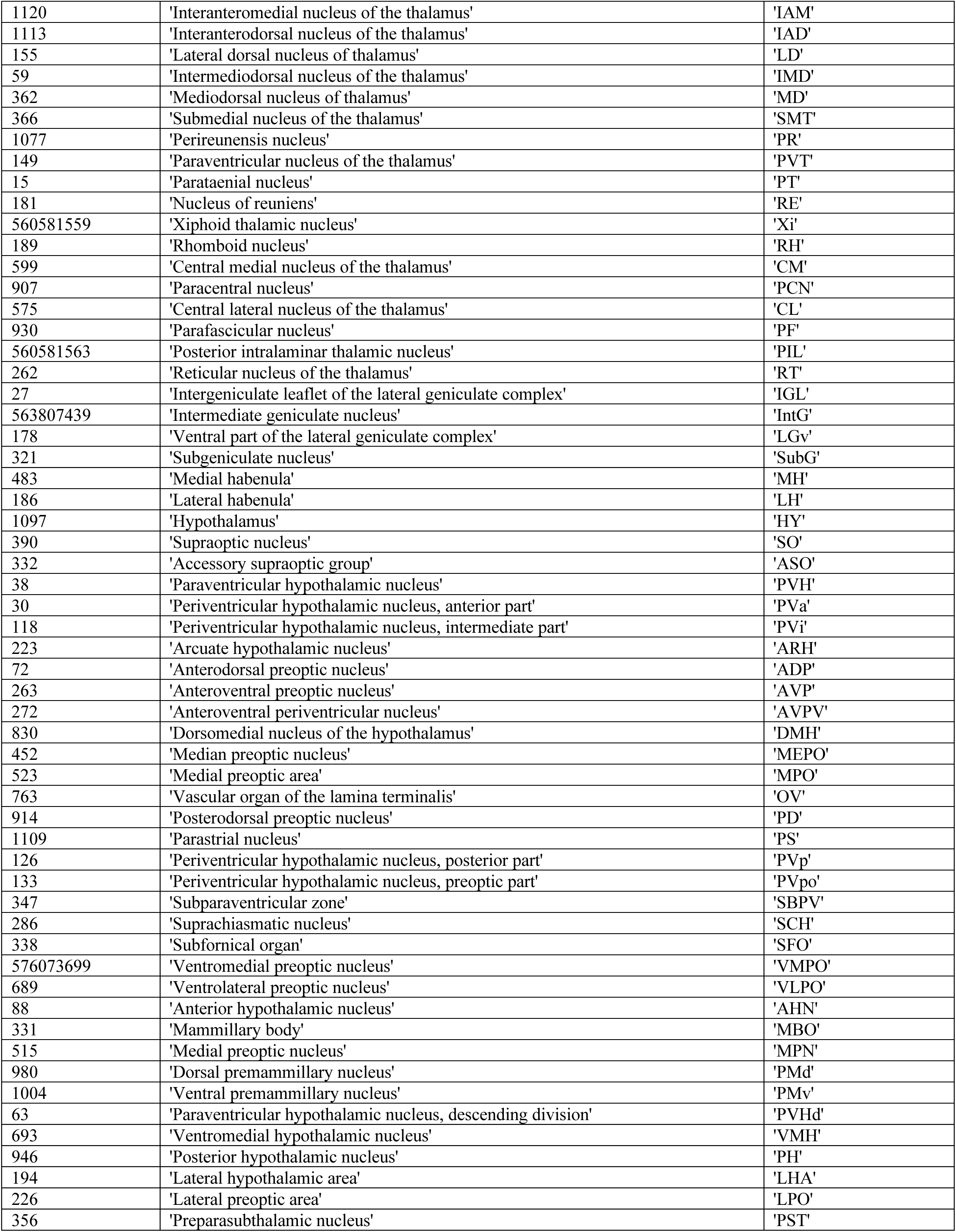
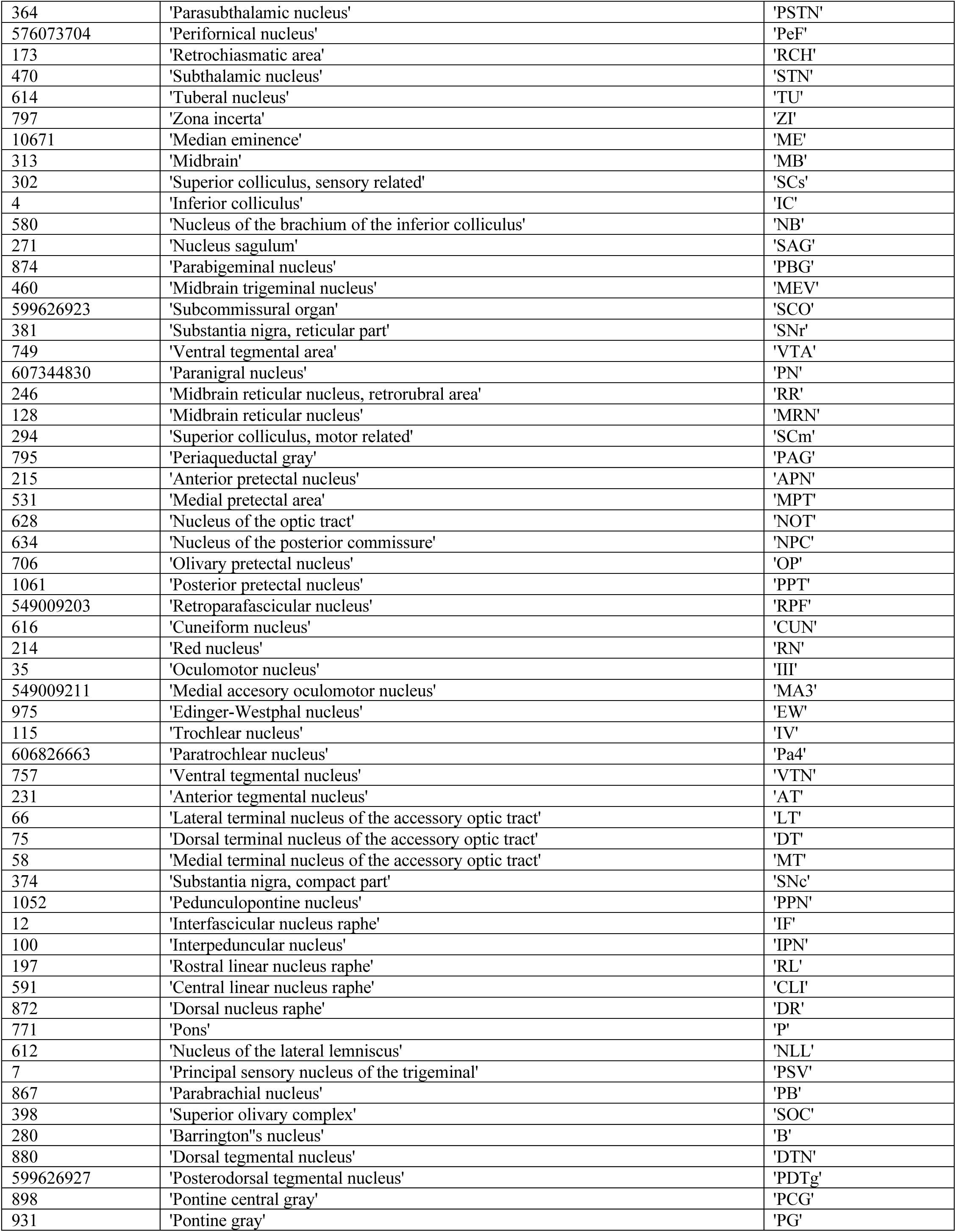
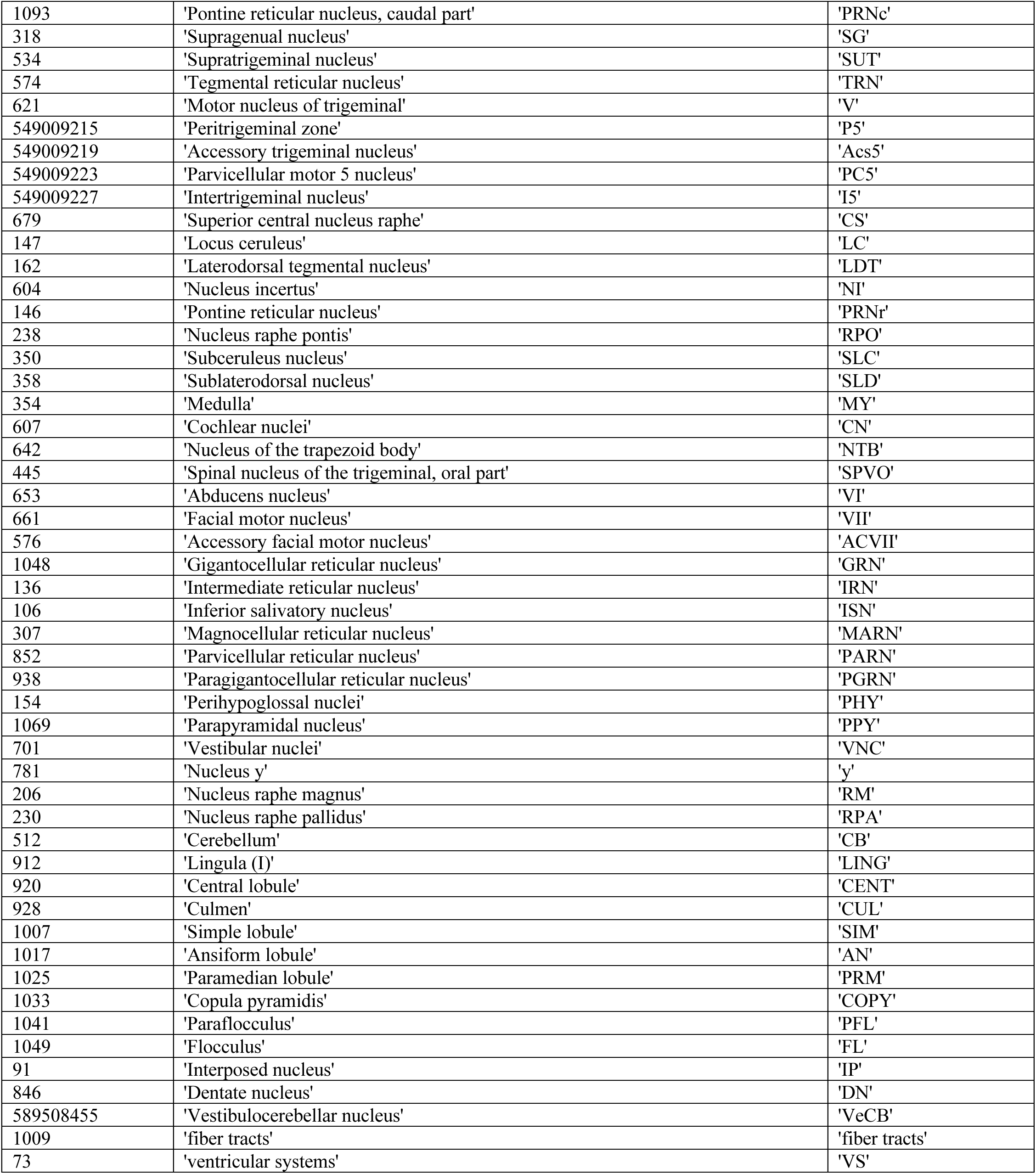
A list of 282 Allen Brain Atlas regions used for Fos map analyses. (see Fig. 3h and Extended Data Fig. 5b)

**Movie S1:** Flythrough of saline-subtracted Fos intensity maps for 9 drugs and the saline group (bottom right).

